# Using Flow Cytometry and Multistage Machine Learning to Discover Label-Free Signatures of Algal Lipid Accumulation

**DOI:** 10.1101/497834

**Authors:** Mohammad Tanhaemami, Elaheh Alizadeh, Claire K. Sanders, Babetta L. Marrone, Brian Munsky

## Abstract

Most applications of flow cytometry or cell sorting rely on the conjugation of fluorescent dyes to specific biomarkers. However, labeled biomarkers are not always available, they can be costly, and they may disrupt natural cell behavior. Label-free quantification based upon machine learning approaches could help correct these issues, but label replacement strategies can be very difficult to discover when applied labels or other modifications in measurements inadvertently modify intrinsic cell properties. Here we demonstrate a new, but simple approach based upon feature selection and linear regression analyses to integrate statistical information collected from both labeled and unlabeled cell populations and to identify models for accurate label-free single-cell quantification. We verify the method’s accuracy to predict lipid content in algal cells (*Picochlorum soloecismus*) during a nitrogen starvation and lipid accumulation time course. Our general approach is expected to improve label-free single-cell analysis for other organisms or pathways, where biomarkers are inconvenient, expensive, or disruptive to downstream cellular processes.

## I. Introduction

There are many biological research tasks for which it is important to measure single-cell behavior [1]. These tasks, which include cell counting, cell sorting, and biomarker detection, are widely conducted using flow cytometry (FCM) [1–3]. Flow cytometry is a high throughput analysis technique that performs rapid multiparametric measurements to inspect and quantify large cell populations and subpopulations [2–9]. FCM analysis is usually conducted by first fluorescently labeling cells, and then quantifying fluorescence intensity of individual cells within large populations. Each cell passes through a laser beam to excite fluorophores, and each cell’s data is recorded by measuring emitted fluorescence intensity at longer wavelengths [5,7,9]. FCM also provides indirect measurements of cell phenotypes through measurements of intrinsic cellular properties, such as cell size and shape by forward-angle light scatter (FSC), and information about cellular granularity and morphology by side-scattered light intensity (SSC) [8,10]. In addition to quantifying cell populations, the related technique of fluorescence-activated cell sorting (FACS) allows researchers to separate cell populations into different subpopulations with respect to their individual properties [8]. As the name implies, sorting decisions are primarily based upon fluorescent labels [1,11].

Despite broad application of fluorescent labels in flow cytometry measurements [10], application of labels can be costly and may require unnecessary effort [12–14]. Biochemical labeling can also alter cell behavior and interfere with cellular processes and downstream analyses by causing activating/inhibitory signal transduction [13,15–19]. Additionally, some biochemical stains require cellular fixation or are toxic, which limits downstream processing when sorting [18,20]. To avoid the adverse effects of biochemical labels, one could engineer synthetic cells to express measurable biomarkers, such as intrinsically fluorescent proteins or luciferases, but such genetic modifications may be costly to develop and may disrupt growth or perturb natural cellular behaviors. A label-free quantification strategy could help prevent these adverse consequences by reducing operation costs and efforts, as well as avoiding side effects of using labels on, or performing genetic modifications to, living cells [12,15]. In label-free quantification of FCM measurements, computational methods are used to quantify targeted cellular information based on measurements from other channels, i.e., from features.

Current label-free quantification strategies employ various methods of machine learning within their analyses to make use of large flow cytometry datasets [12,13,15,17,21,22]. However, in these strategies, the best intrinsic cellular features have been selected based solely on information collected from *fluorescently labeled* cells (for instance, see [12,21]). For some biological processes, if labels indirectly affect intrinsic cell properties within training populations, then these interactions could result in unexpectedly poor quantification of cell populations when tested on unlabeled cells. We hypothesize that FCM datasets could be used to develop label-free quantification strategies *even when signatures are weak and are perturbed* during the training process. In this work, we test our hypothesis by combining supervised machine learning algorithms with analysis of the distributions of single-cell data and their corresponding fluctuation fingerprints [23].

To demonstrate our approach, we conduct feature selection and regression analysis to find optimized label-free feature combinations and quantify lipid accumulation in microalgae cells, that can usually produce lipid content of 15% to 35% (potentially up to 80%), depending upon cultivation conditions, growth media, and algal species [24–27]. For such microalgae to become sources of alternative fuels, it will be necessary to monitor and maximize their ability to accumulate lipids [28]. To enable such quantification, we collect and examine FCM measurements of *Picochlorum soloecismus* under nitrogen replete conditions, and nitrogen deplete conditions that will stress cells and induce them to accumulate lipids. To measure lipid accumulation, we started with a traditional label-based strategy using BODIPY 505/515 fluorescent dye. We measured cell properties with and without the BODIPY stain, and we sought to find signatures in the latter preparation that are capable of reproducing quantities of the former preparation. Using these labeled and unlabeled data, we applied linear and nonlinear supervised machine learning algorithms to select the most informative features and predict lipid content. As opposed to current methods [12,13,15,17,21,22], we show that accurate label-free cell quantification requires rigorous incorporation of statistical information from biological experiments using both labeled and label-free measurements.

## II. Results

Figure 1 depicts our initial strategy for label-free quantification. The first step in our analysis is to grow and measure labeled and unlabeled cell populations as follows:

**Figure 1:**
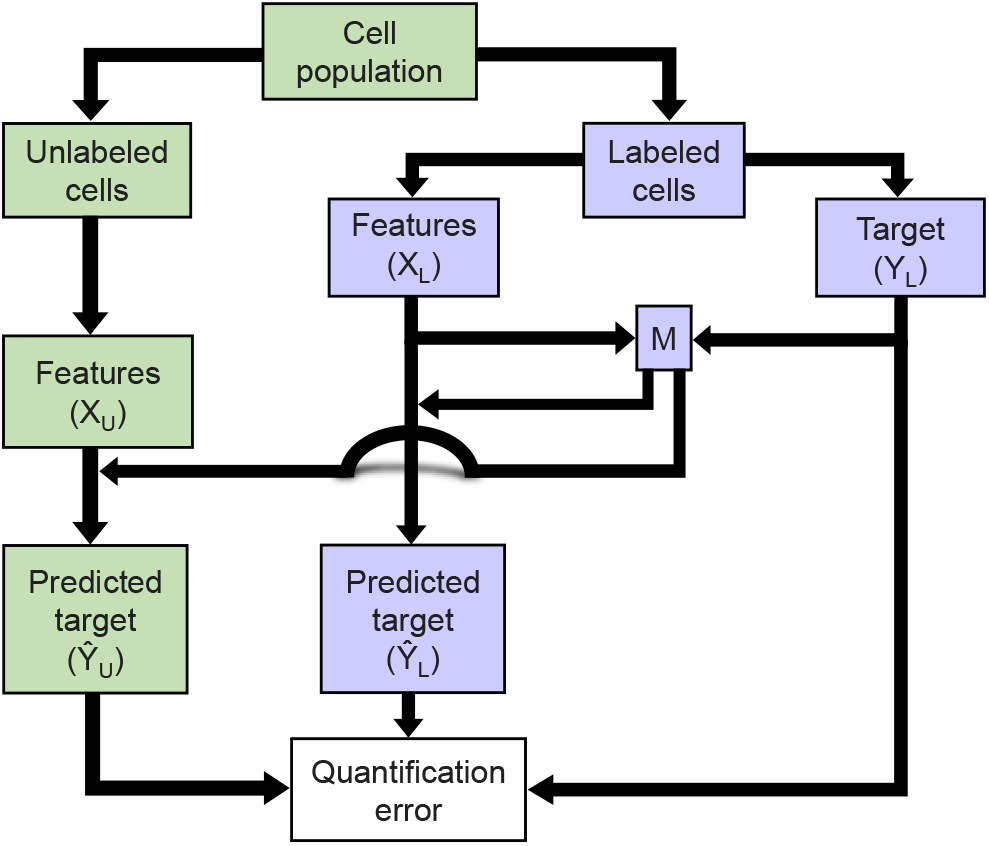
Flow diagram of preliminary regression analysis to quantify lipid content based using intrinsic (presumably label-free) features. The model is learned using labeled data and then tested on both labeled and unlabeled data.

### A. Cell preparation and flow cytometry measurements

*Picochlorum soloecismus* was grown in f/2 media containing half the recipe nitrogen and using Instant Ocean sea salt (Blacksburg, VA) at 38 g/L [29,30]. Cultures were grown at room temperature on a 16 hour light/8 hour dark cycle and mixed by stirring. PH was maintained at 8.25 with on-demand CO_2_ injection when the pH increased above the set-point. Cells were monitored for a total of 46 days following nitrogen starvation. At each time point, we created two identical subsamples as depicted at the top of figure 1. Cells were collected at 23 different days and stored at 4 °C prior to analysis.

To obtain ground truth values for lipid accumulations, we labeled cells in one sub-sample and left the other sub-sample unlabeled. Stained populations of cells were incubated with 22.6 µM BODIPY 505/515 (Thermo Fisher Scientific) with 2.8% DMSO in media for 30 minutes at room temperature prior to analysis. Figure 2 shows representative images of high and low lipid cells. Analysis was conducted using a BD Accuri™ C6 flow cytometer with BD CSampler™ (BD Biosciences). Unstained samples were collected with a set volume of 10 µl on a high flow rate (66 µl/min). For stained samples 10,000 events were collected on a low flow rate (14 µl/min). FCM analyses recorded 13 features per cell, including the 488 nm excitation, 530/30 nm collection channel (FL1) corresponding to the BODIPY dye as well as flow cytometry measurements of forward scatter (FSC), side scatter (SSC) and other fluorescence wavelengths (FL2: 488 nm excitation 585/40 nm collection, FL3: 488 nm excitation 670LP (long pass) collection, and FL4: 640 nm excitation 675/25 nm collection). For the forward scatter, the side-scatter, and each channel FL1-FL4, the flow cytometer measures a pulse of light as each cell traverses the laser beam. Both the height (-H) and the integrated area (-A) of these pulses were collected, providing two measures per channel, per cell.

**Figure 2:**
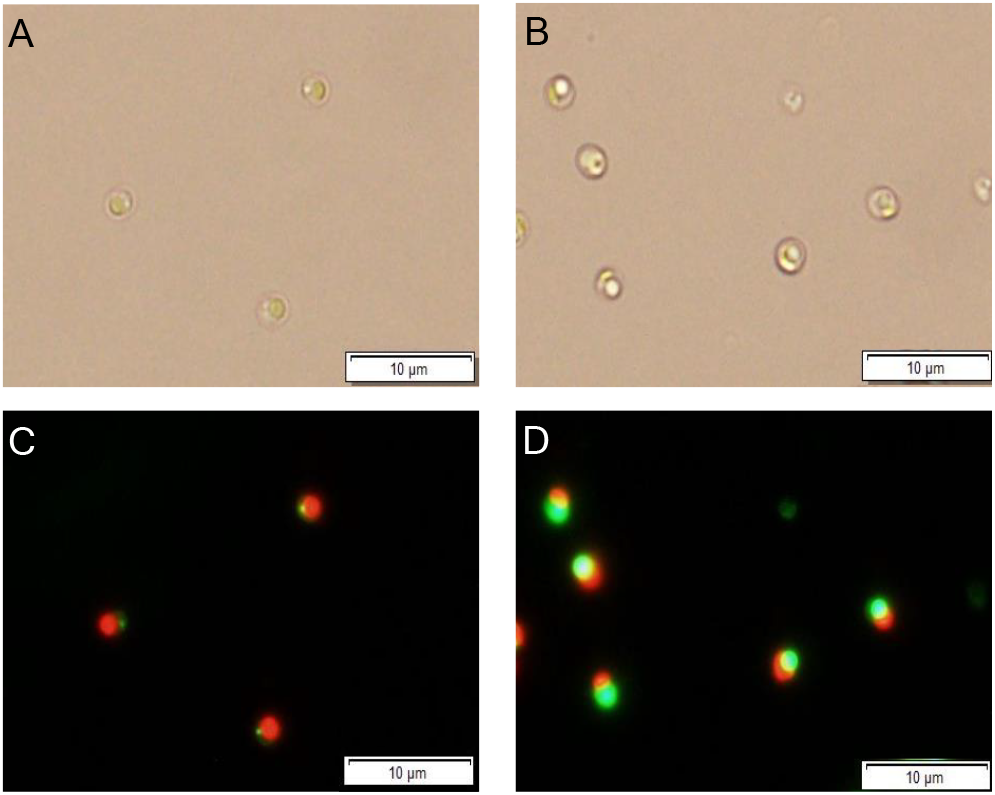
Images of low lipid (A,C) and high lipid (B,D) cells. Panels A and B show the bright field and C and D are overlays of BODIPY staining (green) and chlorophyll fluorescence (red).

All data was exported in .csv format. With these data, we next examined several iterative training-validation strategies to discover signatures within the label-free data that could reproduce the lipid accumulation at all times.

### B. Linear regression analysis

In an initial attempt to identify label-free signatures of lipid content, we considered linear regression applied to match intrinsic features of labeled cells to lipid content (figure 1). In regression analysis, there are two main types of variables: the response variable (denoted *y*) and the explanatory variables (the set of predictors, denoted **x**) [31]. In this study, the response vector is the accumulation of the lipid content for each cell (called the target) and the predictor is a matrix containing the data for intrinsic cellular properties measured by FSC, SSC, and other fluorescence wavelengths (called the features). In regression analysis, the response is approximated as a function of the predictors as

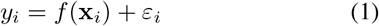

where **x**_*i*_ = (*x*_1_, …, *x*_*N*_)_*i*_ is the vector of *N* intrinsic features for the *i*^th^ cell, and *ε*_*i*_ is a random measurement error for that cell [32]. In linear regression, the response (target) and predictor (feature) variables are assumed to satisfy the linear relationship [32]

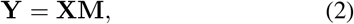

where the vector 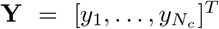 is the vector of targets for *N*_*c*_ training cells; 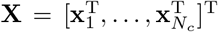 is the corresponding matrix of features for the same cells; and **M** is the regression parameter or the regression coefficient.

Linear regression provides a preliminary insight about potential relationships between the predictor and the response variables. After defining the features and the target, the regression coefficient that minimizes the sum of squared difference of 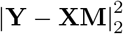 can be calculated using the left inverse of the features matrix, **X**^*−L*^, as follows:

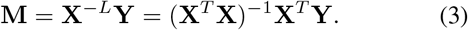

To perform a preliminary regression analysis, we first selected three *training* time points, corresponding to the lowest, the middle, and the highest BODIPY fluorescence intensities (in this experiment, days 1, 14, and 46, respectively). We chose these days to capture the greatest possible range of lipid accumulation phenotypes. For each time point, we considered FCM measurements from a random set of 3000 labeled cells. We computed the regression coefficient, M, by Eq. (3) using the labeled data sets 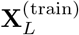 and 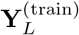. Next, we selected another three *validation* time points, corresponding to the second lowest, another middle, and the second highest BODIPY fluorescence intensities (in this experiment, days 0, 15, and 37, respectively). This time, we extracted information for both labeled, 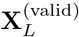 and 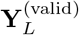, and unlabeled cells, 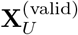. Using the regression coefficient **M** computed previously from training data, we proceeded to predict the lipid content of the labeled and unlabeled validation data sets.

Figure 3 shows the results of applying the simple linear regression analysis using non-label channels from labeled cell preparations in the training phase. Figure 3(A) suggests that the preliminary regression analysis provides a strong estimate for the lipid content using non-label measurements. However, the same regression model failed drastically when those same channels were used to estimate the lipid content in the true absence of labels, and figure 3(B) shows that the difference between predicted and measured values of the lipid content for unlabeled cells is extreme.

**Figure 3:**
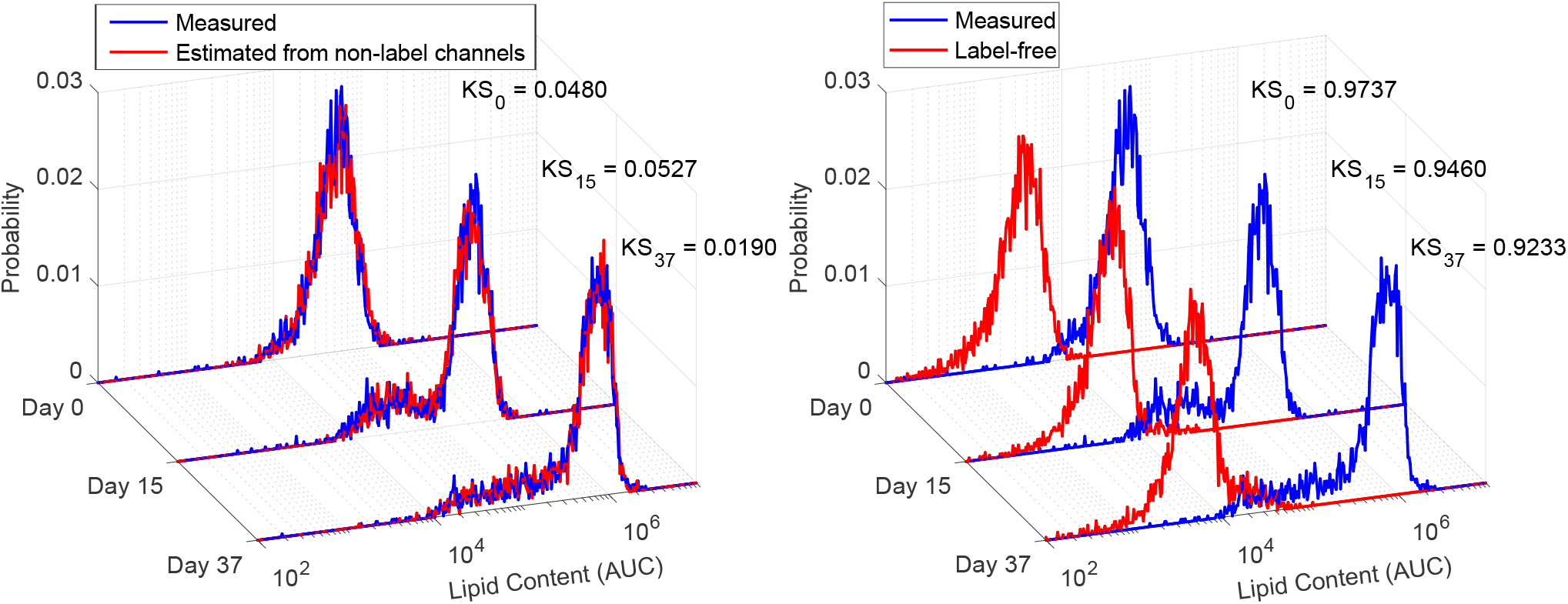
Preliminary regression analysis. (A) Histograms of the lipid content for labeled validation data. The measured lipid contents are shown in blue. The predicted lipid content, estimated from non-label channels, are shown in red. Kolmogorov-Smirnov distances between the distributions are shown. (B) Histograms of the lipid content for unlabeled validation data. The data correspond to days 0, 15, and 37 after nitrogen starvation. All lipid contents are in arbitrary units of concentration (AUC). Bin sizes vary logarithmically.

To explore the similarity of the predicted and measured lipid distributions, we require a metric to quantify the difference between the distributions for lipid estimates and the measured lipid content. Since direct measurement of lipid content is unavailable for unlabeled cells, direct quantification of label-free lipid prediction errors (e.g., single-cell correlation coefficients) is not possible. However, since the labeled and unlabeled cells were sampled from the same original population and at the same time, we reasoned that the labeled and unlabeled populations should have the same distributions or statistics for their single-cell lipid levels. Therefore, to validate label-free predictions, we compare label-free distribution predictions to the labeled measurement distributions using the Kolmogorov-Smirnov statistic (KS). In one dimension, the KS ranges from zero (perfect agreement) to one (total disagreement) and can be defined independently of scaling or units to discern the difference between two distributions [33,34]. We found that KS distances between non-label channel predictions and measurements for labeled cells were very small (0.0480, 0.0527, and 0.0190 for the three validation time points), but the corresponding KS values for the label-free predictions were almost completely incorrect (0.97, 0.95 and 0.92 for the same time points). Extended results for the linear regression are provided in supplementary Figs. S1 and S2.

### C. Nonlinear approaches

To generalize our initial simple linear regression approach, we then added new features corresponding to all possible products of the individual features as follows:

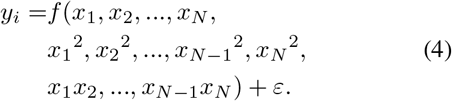

This expanded linear regression analysis, which uses all possible quadratic features, is referred to as the *quadratic* regression model. To further generalize the analysis, we also formulated a multilayer perceptron neural network (MLPNN) [35] and also applied the gradient boosting machine learning (GBML) method presented by Blasi et al. [12] to predict the BODIPY signals in our FCM measurements (see sections 2 and 3 in the supplementary information for details on MLPNN and GBML). However, as shown in supplementary figures S3-S5, each of these advanced approaches appeared to work very well on the *labeled* training and validation data, but all were insufficient to predict the lipid content for *unlabeled* data.

### D. Feature selection

To explain the failure of the labeled-cells-trained regression model on unlabeled cells, we suspected that some channels in the flow cytometer might be adversely affected by application of the BODIPY stain. Indeed, after analyzing the labeled and unlabeled features, we observed that some intrinsic features (FL2-A and FL2-H, corresponding to the second channel of the flow cytometer) change substantially when BODIPY is added to the cells (supplementary figure S6). This channel is the closest to the FL1 channel that measures the lipid content, where the BODIPY fluorescent dye is added. Moreover, it is conceivable that the level of this disruption could be correlated with the amount of lipid in the cells, which means that it could be equally present in both training and validation data for the labeled cells. As a result, these changes could disrupt the training and cross-validation procedures and account for prediction failure when tested on unlabeled cells.

To mitigate this effect, we removed features FL2-A and FL2-H from the regression analysis and then repeated the linear regression. We found that removing these disrupted features led to substantial improvement for the quantification of unlabeled data (supplementary figure S7: KS improved from 0.92-0.97 in figure 3 to 0.11-0.38 in figure S7(B)). The supplementary figure S8 provides extended plots of the outcomes of regression analyses upon removal of disrupted features. It is interesting to note that removal of disrupted features reduces accuracy of lipid prediction for labeled cells. This occurs because the labeling inadvertently modulates some “intrinsic” features in the labeled cells and introduces extraneous feature-target correlations that are actually detrimental to predictions for unlabeled cells. A troublesome consequence of these correlations between labels and intrinsic features is that these disrupted features are immune to removal when cross-validation analysis is applied exclusively to labeled cells.

Next, we used the genetic algorithm on combinations of labeled and completely unlabeled data to explore if further feature reduction could enhance label-free classification. The supplementary figure S7(C,D) shows the results following the application of the genetic algorithm, which automatically selected FSC-A, SSC-A, FL3-A, FSC-H, and the width of the signal as the most informative features. Down-selecting to these most informative features resulted in a slightly smaller KS distance (0.10 - 0.35) between measured and predicted values of the lipid content for unlabeled cells. Extended results are provided in supplementary figure S9.

During automated feature selection for linear regression (supplementary figure S7(C,D)), we did not incorporate higher order effects (e.g., “interactions”) between predictor variables. To enhance our modeling and potentially extract more information from the data, we added an expanded set of products of feature values to the input. Expansion of the input matrix of features to include quadratic and first order interaction terms, followed by label-free feature selection via the genetic algorithm, resulted in a slight improvement to label-free predictions for the lipid content. For more detailed results after introducing the quadratic features and application of the genetic algorithm on higher order effects, see figures S7(E,F) and S10 in the supplementary information. In this case, the genetic algorithm identified the product of FSC-A and FL4-H, the square of FSC-H, and the product of FL4-H and signal width as the most informative attributes. Selected features by the genetic algorithm on linear and quadratic features are presented in more detail in supplementary Table S1.

### E. Weighted model

After cross-validation and feature reduction, the predictions using label-free data had improved substantially, but we noticed that there remained some substantial systematic prediction errors. In particular, predictions using a single regression model appeared to be biased toward the average lipid levels and led to over-prediction of low lipid populations (early time points) and under-prediction of high lipid content populations (late time points). We hypothesized that this bias to the middle could be corrected by allowing the model itself to adapt in accordance with signatures in the label-free data.

To test this idea, we introduced a new strategy based on weighted models that could be learned from all measurement of unlabeled features. To achieve this weighted model, we first learned three separate regression coefficients **M**_1_, **M**_2_, and **M**_3_ based on the three training time points (days 1, 14, and 46). These models were fixed for all subsequent computations. For any arbitrary population, a new combination model could be formulated as a weighted sum:

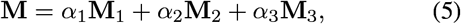

where the weights **a** = [*α*_1_, *α*_2_, *α*_3_] would be specific to any new population of unlabeled cells.

We then sought to learn a secondary model to estimate **a** from populations of unlabeled data. For this task, we defined 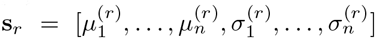 as a vector that contains the population means and standard deviations of each feature (including quadratic features) in any population of unlabeled cells. It is important to note that because the unlabeled cells are not treated with BODIPY, the statistics contained in **s**_*r*_ can include the 530/30 nm channel, which allows access to previously unutilized information in the unlabeled cells. We then constructed the population sample statistics matrix 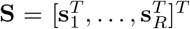 using *R* different randomly sampled sub-population from the original training and validation data. For each *r*^th^ random population, we also performed a computational search to find an optimized model scaling factor **a**_*r*_ that yields the best possible comparison between measured and predicted targets in the training and validation data, and we collected these into the matrix 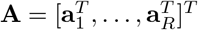.

With these definitions, we formulated a secondary regression analysis for **a**_*r*_ as a function of **s**_*r*_ with the assumed linear form

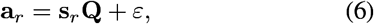

for which we could estimate the weight quotient **Q** as

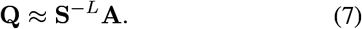

In this expression, **Q** defines a relationship between the unlabeled features (from computing **s**) and the weights (**a**). To prevent over-fitting in the determination of the weights, we generated another set of random population samples from our training and validation data, and we used the genetic algorithm to down select among the best columns of **S** (or rows of **Q**) to utilize for the estimate of **a**. Once fixed using the training and validation data, the multi-scale regression operators **M**_1_, **M**_2_, **M**_3_ and **Q** could be applied to any new data sets **X**_*U*_ and their summary statistics **s** to calculate **a** = **sQ**, estimate **M** using Eq. (5), and predict the lipid content using Eq. (2).

Figure 4 shows the results of our new weighted-model label-free quantification strategy for labeled cells (figure 4(A)) and unlabeled cells (figure 4(B)). It can be seen here that using model weights, which are based only on statistics of unlabeled features, enables the model to predict the BODIPY signal with a remarkably high accuracy. The expanded weighted model analysis allows for a substantially improved ability to quantify lipid content for both labeled and unlabeled cells. The very small KS distance (0.14, 0.09, and 0.09) on the three validation time points represent an exceptional success in predicting the BODIPY signals based on label-free measurements.

**Figure 4:**
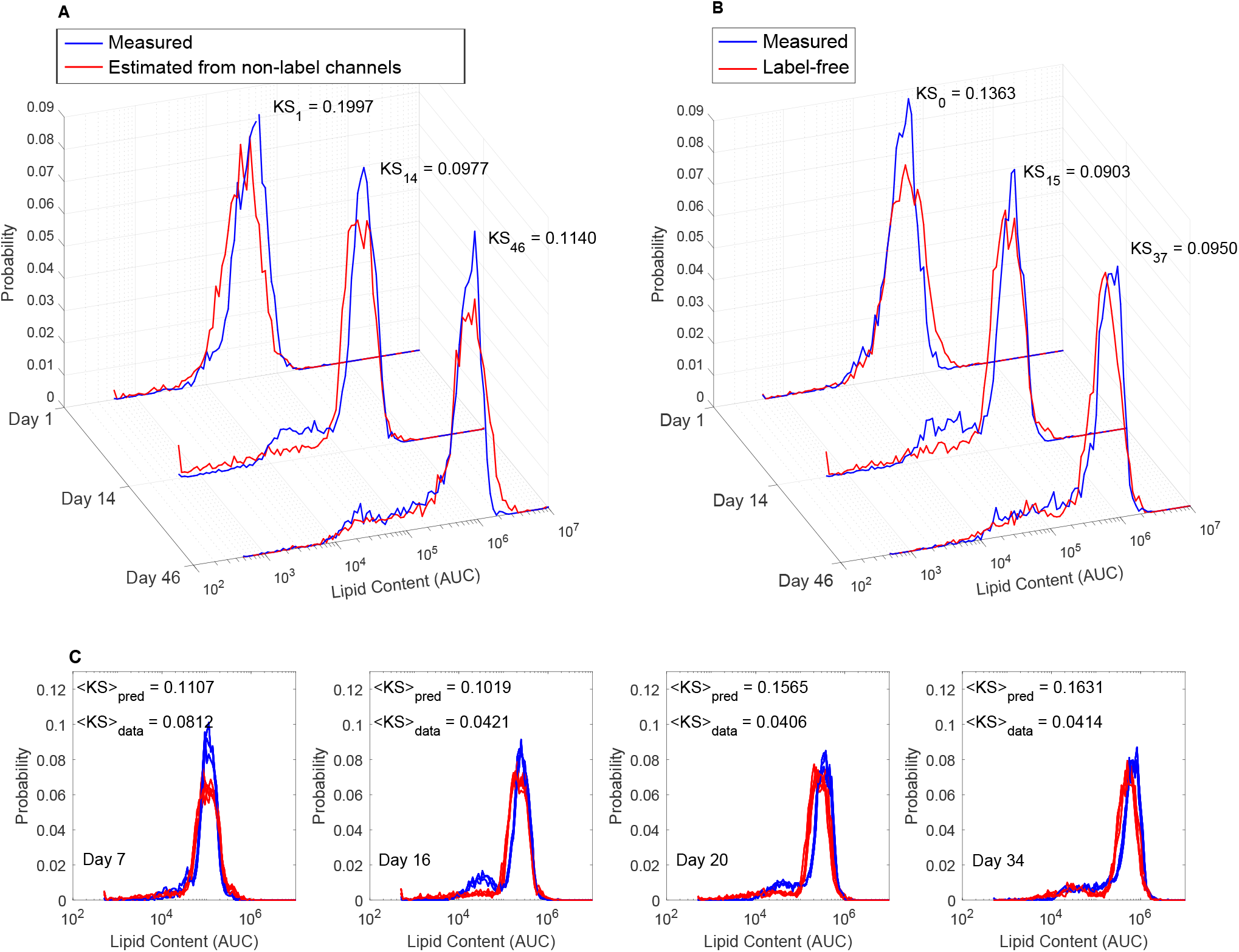
Results of the weighted model. Distributions of lipid content for (A) labeled training data, and (B) unlabeled validation data. KS distances between distributions are shown. (C) Testing the final strategy on four unlabeled testing time points: Days 7, 16, 20, and 34. See supplementary figure S10 for corresponding results for all 17 testing time points. “KS data” is the average KS distance between measured lipid distributions. All lipid contents are in arbitrary units of concentration (AUC).

### F. Testing the final model on new time points

After we validated the final label-free lipid estimation model, we fixed all parameters and sought to test it for label-free quantification in new circumstances. Figure 4(C) shows that the final model yielded exceptional prediction accuracy of the BODIPY signal for this previously unseen testing data at time points corresponding to days 7, 16, 20 and 34, and supplementary figure S12 shows the corresponding predictions at the remaining 17 time points not used in the training data set. Figure 5(A) also shows that the trained model correctly quantified average and standard deviation of lipid accumulation (in log scale) at each day following nitrogen starvation. We note that the training and validation data were taken at only three time points each, yet the model sufficed to predict the lipid levels for all of the remaining 17 time points. Figure 5(B) shows the changes in model weights, **a**, which were estimated solely from the statistics of the unlabeled data (three biological replicas per time point) and without any information about the time of measurement. The figure demonstrates that the secondary model correctly adapts these weights from a domination of *α*_1_ at early times, *α*_2_ at middle times and *α*_3_ at late times.

**Figure 5:**
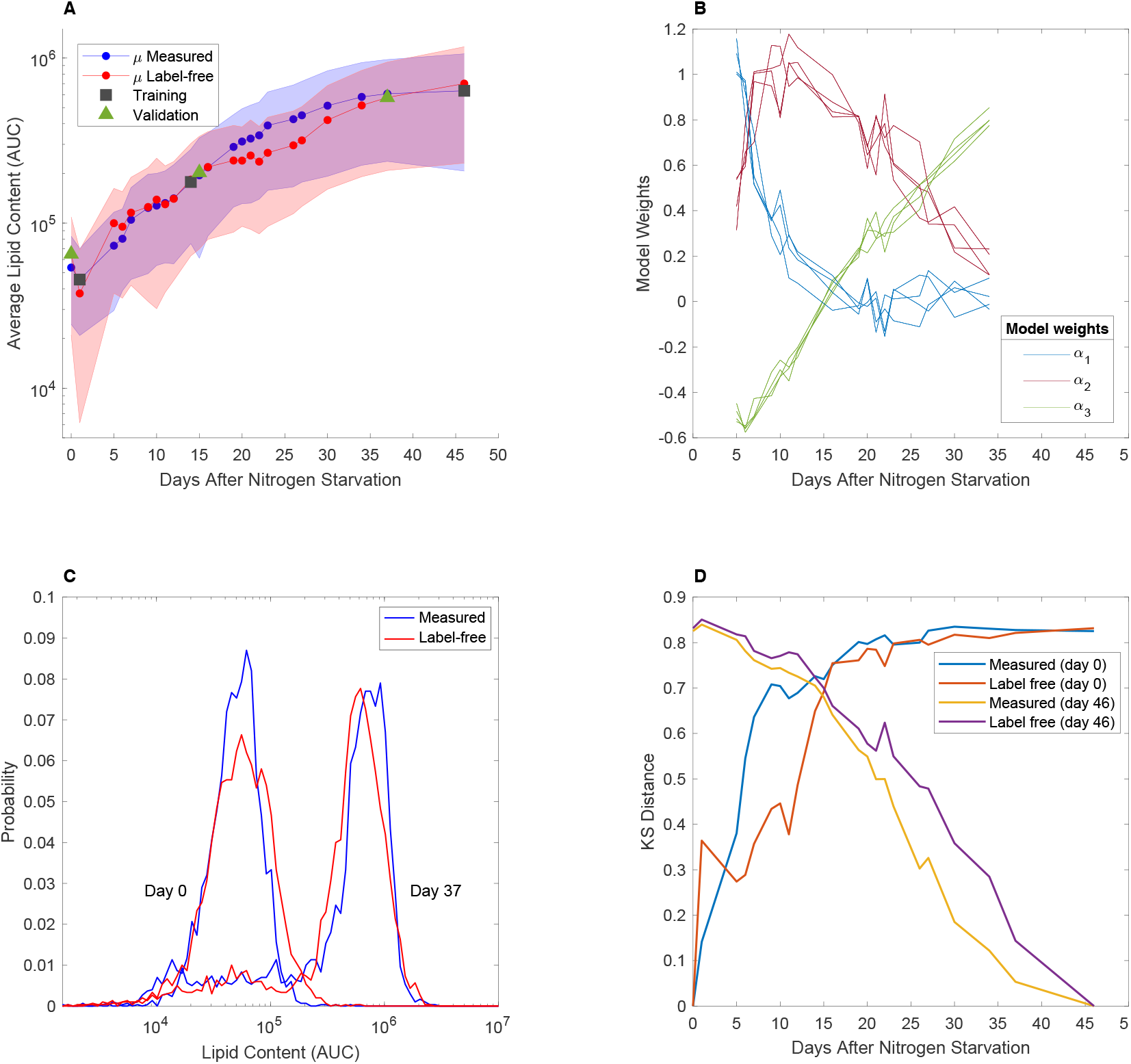
Results of the weighted model. (A) Average lipid content at each day after nitrogen starvation. The blue and red shaded areas show the standard deviation as measured and predicted, respectively. Training, validation, and testing time points are shown in black rectangles, green triangles, and red and blue circles, respectively. (B) Model weights calculated based on label-free information of the testing data at each day after nitrogen starvation. *α*_1_,*α*_2_, and *α*_3_ correspond to the weights for model 1, model 2, and model 3, respectively. (C) Comparison of the distributions of the lipid content between days 0 and 37 after nitrogen starvation. The KS distance between the measured values is 0.8277. The KS distance between the label-free values is 0.8213. (D) Change in the KS distance with respect to time. Blue and red show the changes in KS distance between day 0 and other days after nitrogen starvation for measured and label-free data, respectively. Yellow and purple show the changes in KS distance between day 46 and other days after nitrogen starvation for measured and label-free data, respectively. All lipid contents are in arbitrary units of concentration (AUC).

Figure 5(C) compares how much the lipid distributions changed over the course of the nitrogen starvation experiment as quantified using labeled (blue) or label-free (red) strategies. We found that the KS difference between the initial and final time points found for the label-based and label-free measurements were in excellent agreement of 0.83 and 0.82 respectively. Using the KS distance, we can now compare the dependence of the population distribution on changes to underlying variables, using analyses similar to those demonstrated in [34] to quantify population responses to external regulatory factors. In our case, figure 5(D) shows the KS distance between distributions at variable time *t* compared to the initial or final times and calculated from the direct lipid measurements (blue, gold) or label-free estimates (red, purple). Once again, we find that the comparisons using label-free measurements are in excellent agreement with the label-based measurements for all time points throughout the nitrogen starvation process.

### G. Relationships of label-free features to lipid content

Table 1 presents the most informative features selected by the genetic algorithm for the construction of the regression analyses at the three training times and for the multimodel weighting coefficients. Tables S1 and S2 provide the specific numerical values for all regression coefficients. In this case, the feature selection results can be interpreted in terms of known biology. To aid in this interpretation, figure 6(A) shows the median levels of three of the most important label-free features (SSC, FL3-A, and FL4-A) and the median measured lipid content versus time after nitrogen starvation. First, we note that the label-free SSCA measurement is positively correlated with lipid content throughout the time course. This is easily explained by noting that SSC is indicative of granularity of the cells, and as lipids generally accumulate in lipid bodies as shown in figure 2, these bodies are likely to account for the increased scattering measurements in the flow cytometer. Second, we note that the fluorescence channels FL3 and FL4 exhibit weak negative correlations to lipid content at later times. Much of the fluorescence measured in these channels is likely to originate from chlorophyll. Our analysis suggests that nutrient deprived cells, which are accumulating lipids as a stress response, slowly deplete their levels of chlorophyll over time, an observation that is consistent with previous studies applying bulk cell culture analyses to other species of algae [36,37]. For the secondary regression analysis used to define the weights of the regression analyses, the optimum found by the genetic algorithm relied primarily on these same features, but were supplemented by statistical information from other fluorescence channels, including the 530/30 nm channel that was discarded to conduct training on labeled cells.

**TABLE 1:**
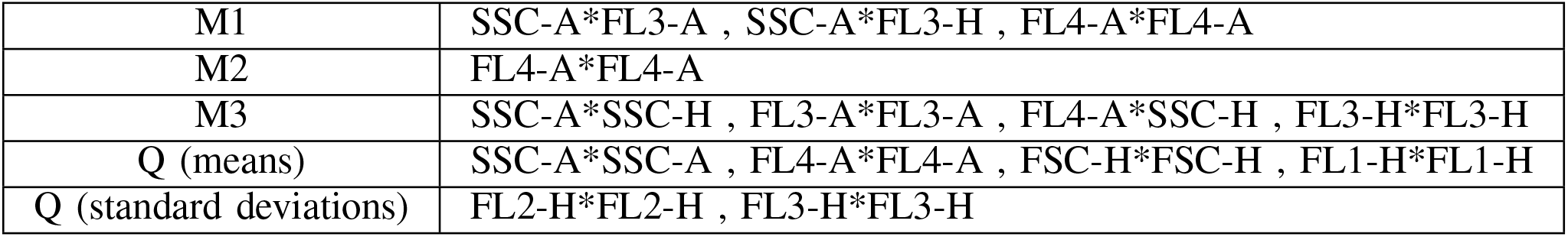
Selected features at the three training time points corresponding to **M_1_**, **M_2_**, and **M_3_**. The genetic algorithm selects feature means, and feature standard deviations for the weight quotient **Q**.

|  |  |
| --- | --- |
| M1 | SSC-A*FL3-A , SSC-A*FL3-H , FL4-A*FL4-A |
| M2 | FL4-A*FL4-A |
| M3 | SSC-A*SSC-H , FL3-A*FL3-A , FL4-A*SSC-H , FL3-H*FL3-H |
| Q (means) | SSC-A*SSC-A , FL4-A*FL4-A , FSC-H*FSC-H , FL1-H*FL1-H |
| Q (standard deviations) | FL2-H*FL2-H , FL3-H*FL3-H |

**Figure 6:**
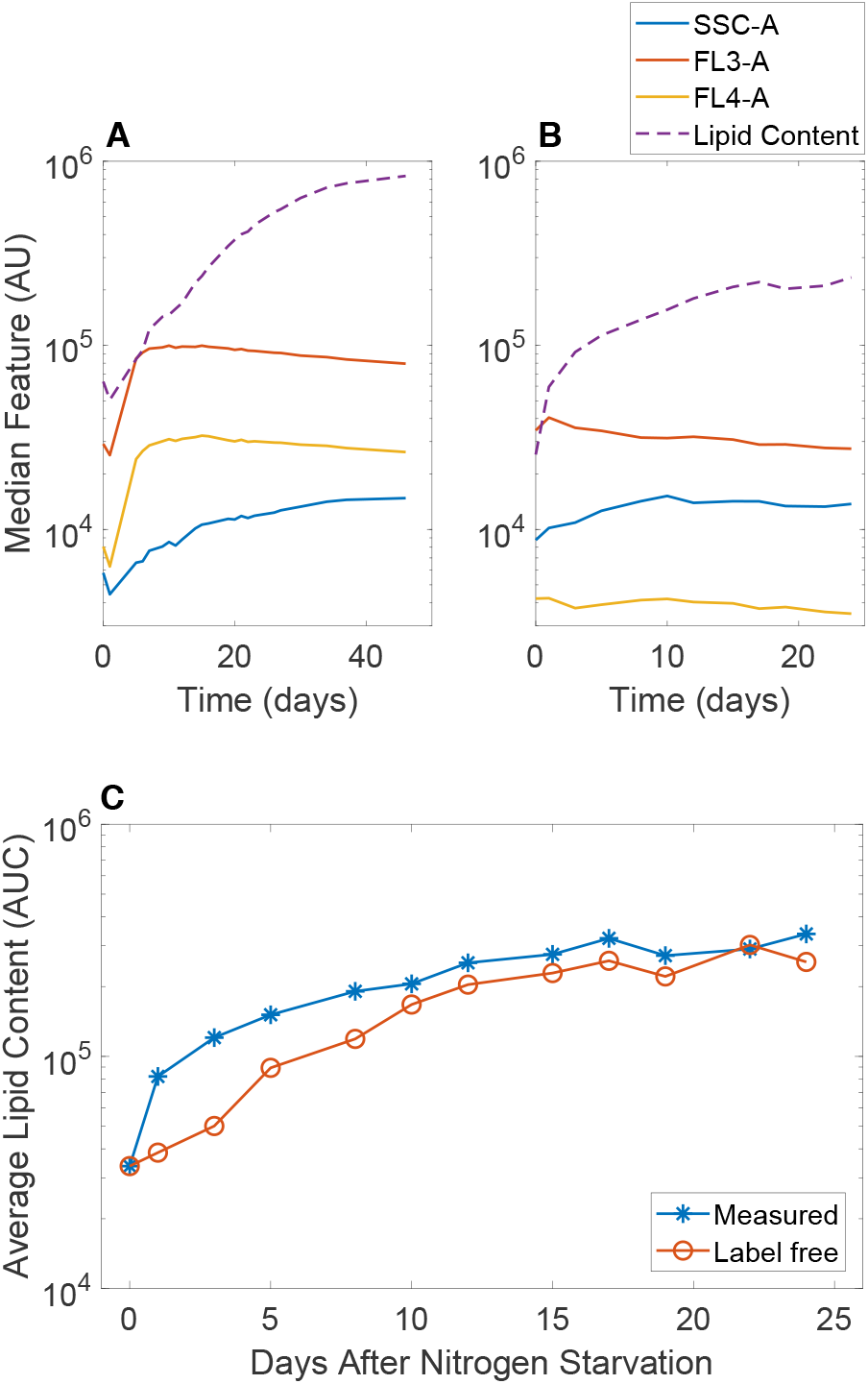
Testing the final model on independent cell populations and with a new flow cytometer. (A,B) The median values of the most informative label-free features (solid lines) and the label-based lipid measurement (dashed lines) versus time after nitrogen depletion using (A) the original cell preparation and (B) for a subsequent independent cell preparation measured with a different flow cytometer. (C) Evaluation of the final weighted model’s estimates of lipid content when applied to new cells with the new flow cytometer. Blue shows the lipid accumulation for the measured values of the lipid content at each day after nitrogen starvation. Red shows the label-free predictions of the lipid content for the same days. Both curves have been normalized relative to the measure lipid level on Day 0. Lipid content are in arbitrary units of concentration (AUC).

### H. Generalizing the model on a different flow cytometer

A substantial impediment to the development of label-free strategies for flow cytometry analysis is that collective dynamics can cause one cell population to behave differently from another, even when they are prepared in similar environmental conditions. Moreover, it is not uncommon that two flow cytometers, with different settings or containing different optical components, could yield different measurements, even when used to measure the same cell populations. With these issues in mind, we next sought to test the generality of our approach when applied to a new preparation of *P. soloecismus* over time during nitrogen starvation and quantified using a different flow cytometer (BD Accuri™ C6 Plus, which has a different fluidics system) with matching detectors. Without any re-training of our previous model (i.e., using the same features and model parameters identified above), we sought to quantify the lipid accumulations over time for the new data set. Owing to variation in the flow cytometer and its settings, the quantitative values for the measurements changed considerably, as can be seen in figures 6(A) and 6(B), which show the median measurements for the label-free features and lipid measurement for the old and new data sets, respectively. Despite substantial differences, figure 6(C) demonstrates that our previous model still correctly captures the trend of increasing lipid accumulations over time based on the label-free information collected using the new cell preparation and new flow cytometer.

### I. Simulating the model’s capability to perform sorting

Finally, we used our weighted model to explore the possibility that it could be used to sort unlabeled cells according to the lipid content within those cells. To simulate this situation, we generated mixed cell populations by combining 2500 cells randomly chosen from the initial (low lipid) time point and 2500 cells randomly chosen from the final (high lipid) time point. We then applied the previously identified model ({**M**_1_, **M**_2_, **M**_3_, **Q**}) on the entire mixed population to predict the distribution of lipids from the label-free features. Figure 7 shows the results of this mixture for the quantification based upon labels (panel A) and based upon label-free measurements (panel B). In each case, the green lines correspond to the subpopulation taken from the early time points, the purple lines correspond to the label-free measurements, and the black lines correspond to the full mixed distributions. We assumed an optimal gate (vertical dashed line in figure 7), and we asked what fraction of cells from the green/purple sub-populations would be correctly assigned to the low/high populations. As a benchmark, we found that the label-based sorting accuracy was 72.84% to classify low lipids cells and 94.32% to classify high lipid cells (figure 7(A)). The label-free sorting accuracy performed equally well at 79.64% correct classification of low lipids and 92.2% correct classification of high lipids and (figure 7(B)). These results suggest that label-free classification could be used in principle for sorting applications, but full evaluation of this hypothesis, as well as strategies to optimize label-free sorting gates, remain to be validated through future experimental investigations.

**Figure 7:**
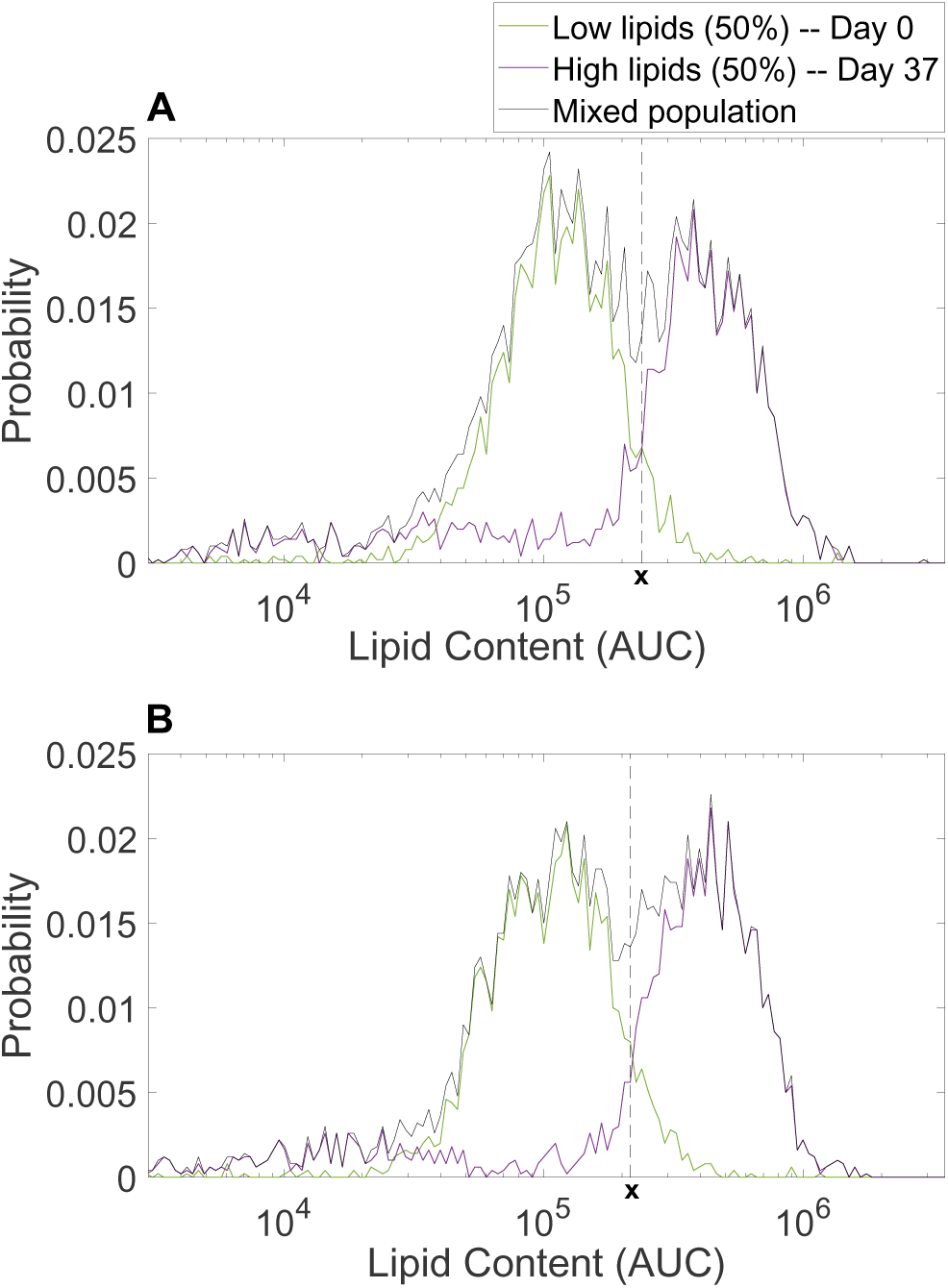
Simulation of a typical cell sorting experiment. (A) Sorting the labeled cells based on our optimized weighted model into high and low lipid content. Label-based sorting accuracy of 94.32%. (B) Sorting the unlabeled cells based on our optimized weighted model into high and low lipid content. Label-free sorting accuracy of 92.2%. For all panels, the lipid content are in arbitrary units of concentration (AUC). Simulated populations were generated by combining experimental data from 2500 cells from Day 0 and 2500 cells from Day 37.

## III. Conclusions

Single-cell quantification and classification are crucial tasks in many biological and biomedical applications, and flow cytometry (FCM) is one of the most common tools used for these tasks. Computational strategies have substantial potential to identify label-free markers and mitigate the expense or disruptive effects of traditional FCM analyses. In this article, we have demonstrated the use of mathematical tools and statistical methods, including regression analysis and machine learning to extract quantitative information from intrinsic properties of unlabeled cell populations. We discovered that computational classifiers that are learned using intrinsic features measured in labeled cell populations may appear to be highly predictive when compared to other labeled cells, but these same models may then fail dramatically when tested on truly label-free data (figures 3 and S2-S5).

The key to our integrated strategy is careful consideration of the variations within heterogeneous single-cell populations. Drawing inspiration from our past work to identify gene regulation models from single-cell distributions [23,38,39], we reasoned that distributions of labeled and unlabeled cell populations should have shared statistics that could help circumvent the issue of data disruption due to label applications. Under that inspiration, we developed a multi-stage regression approach that incorporates collections of both labeled and unlabeled data in the same conditions. From these data sets, we learn which features’ statistics are conserved, which features vary between different treatments, and which features are most valuable to predict lipid content in unlabeled cells when trained using labeled cells. Figure 8 depicts a flow diagram of our new approach and its three main components of (i) linear regression applied to features and feature products to discover the correlations between intrinsic features and lipid content within labeled cells; (ii) genetic algorithms to automatically select features that contain useful information, but which avoid misleading or distracting artifacts contained within large FCM datasets; and (iii) a new model-weighting strategy to allow application of different statistical models in different situations.

**Figure 8:**
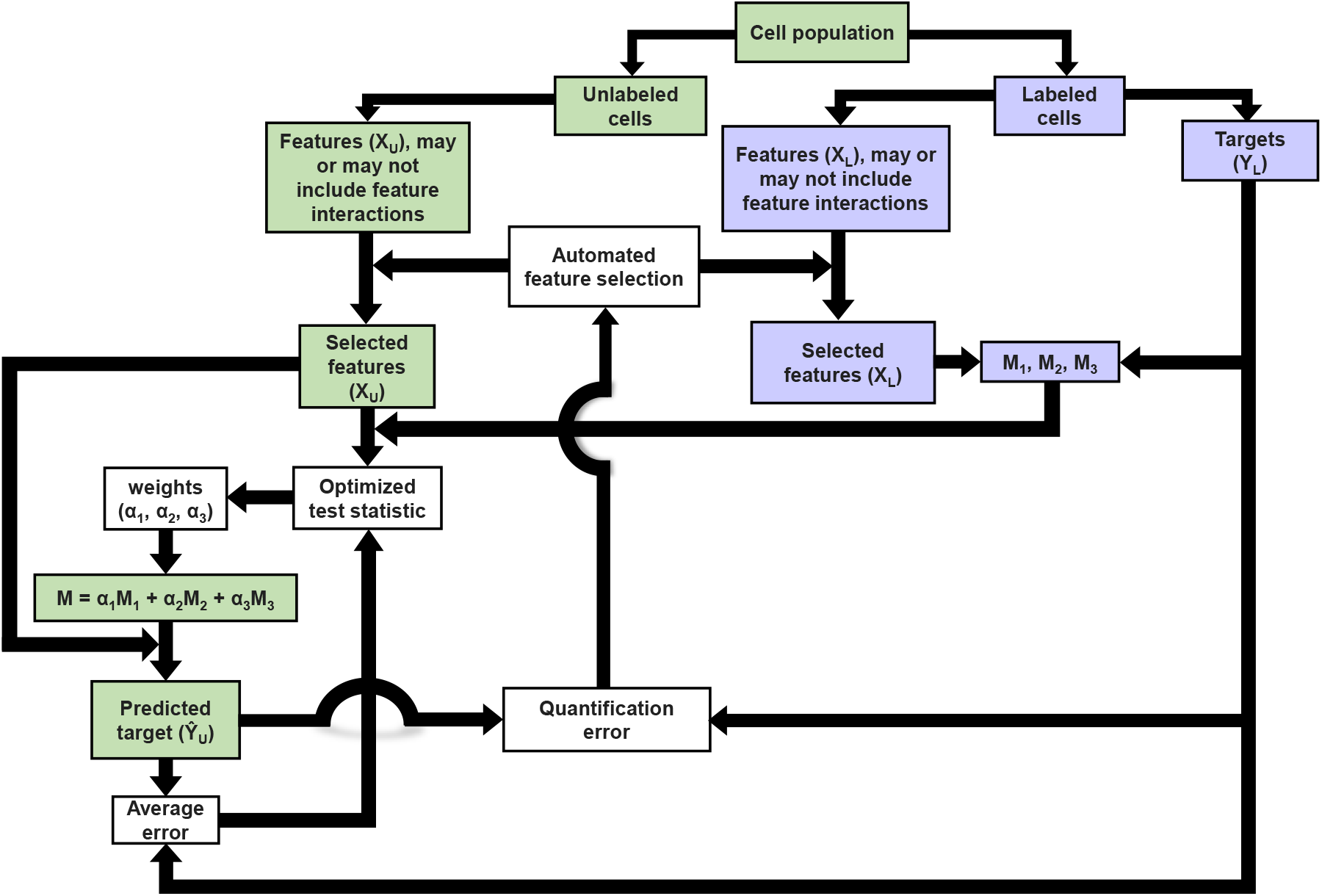
Flow diagram of the final multi-stage label-free quantification strategy.

The combination of regression analyses, genetic algorithms and model weighting approaches yields a final set of models and weights that are uniquely determined from the statistical properties of unlabeled cell population measurements. Using this approach, we can then extract sufficient information to provide efficient label-free quantification of lipid content in *Picochlorum soloecismus* over time during nitrogen starvation. Our final model accurately estimates lipid content distributions over time that span several orders of magnitude (figures 4, 5, and S12). Moreover, although direct verification of lipid content for unlabeled single-cells is not possible, our final regression models preserved single-cell prediction accuracy for lipid content in labeled cells, especially at later time points when lipid content is highest (Pearson’s correlation coefficient of R ≈ 0.74-0.87; see supplementary figure S11).

Together, the proposed computational tools could help circumvent the need for biochemical labels to reduce expense, simplify sample preparation protocols, and open new avenues for single-cell research. For example, label-free quantification will be instrumental to sort cells into different subpopulations, without the (potentially terminal) cellular disruptions associated with standard biochemical markers or with genetic modifications needed to express fluorescent proteins. Once trained through several rounds of regression and genetic algorithms, our final model for algal lipid quantification reduces down to a simple linear operation applied to a handful of seven second-order products of features of the unlabeled cells. Such operations are easily computed in less than a microsecond per cell, making the label-free analysis ideal for use in gating and sorting applications as a stand-in for fluorescence in fluorescence-activated cell sorting (FACS) analyses.

Applied to algae cultivation for biofuels and bioproducts (food and feed ingredients), real time monitoring of cultures can provide information on the health and productivity of the algal cells. This allows for harvesting when the cells are at maximum yield, or prior to being overtaken by pests or predators. Moreover, the ability to monitor without the addition of dyes increases the speed of analysis and decreases costs. Additionally, the ability to sort cells of interest without labels would enable selection of subpopulations with a desired phenotype of interest, such as higher content of lipid or other value added products, such as specialty oils or cannabinoids. This type of selection would allow for directed improvement of strains without direct genetic modification.

Finally, the ability to analyze and sort cells without labels is broadly useful as a strategy to improve productivity of cell cultivation operations for a variety of applications in the medical or pharmaceutical industry. While our machine learning approach wont substitute for the use of labels in every application in flow cytometry, it is applicable in cases where there are subtle morphological features that accompany a biological change in a cell. This approach allows identification of those morphological changes that would be normally detected with a label, and in this case we were able to identify them with machine learning and allow detection in the absence of a dye.

## IV. Acknowledgments

Research reported in this publication was supported by the National Institute of General Medical Sciences of the National Institutes of Health under award number R35GM124747 and by a Laboratory Directed Research and Development grant from Los Alamos National Laboratory (project 20130239ER).

