## Supplemental Information for "Using Flow Cytometry and Multistage Machine Learning to Discover Label-Free Signatures of Algal Lipid Accumulation"

### Supplementary Information

Mohammad Tanhaemami

#### Contents

**1 Flow Cytometry Measurements and Data Acquisition** **2**

**2 Gradient Boosting Machine Learning (GBML) Suggested by Blasi et al.** **2**

**3 Multilayer Perceptron Neural Network (MLPNN)** **2**

### 1 Flow Cytometry Measurements and Data Acquisition

After preparing samples of labeled and unlabeled *Picochlorum soloecismus*, we conducted measurements using a BD Accuri™ C6 flow cytometer (BD Biosciences). The flow cytometer contains the following channels:

- FSC Forward scatter (low angle scatter, generally related to size)
- SSC Side scatter (90-degree scatter, generally related to granularity)
- FL1 488nm excitation, 530/30 collection, this is the channel we use to look at BODIPY
- FL2 488nm excitation, 585/40 collection
- FL3 488nm excitation, 670LP (long pass) collection, this is the channel we use to look at auto fluorescence
- FL4 640nm excitation, 675/25 collection, auto fluorescence is also strong here

As the cell transverses the laser beam, a pulse is created for which "H" is the height, "A" is the area, and "Width" is the width of that pulse.
The table below presents the information provided at each channel in our flow cytometer.

| Feature 1 | Feature 2 | Feature 3 | Feature 4 | Feature 5 | Feature 6 | Feature 7 |
|---|---|---|---|---|---|---|
| 'FSC-A' | 'SSC-A' | 'FL1-A' | 'FL2-A' | 'FL3-A' | FL4-A' | 'FSC-H' |
| Feature 8 | Feature 9 | Feature 10 | Feature 11 | Feature 12 | Feature 13 | |
| 'SSC-H' | 'FL1-H' | 'FL2-H' | 'FL3-H' | 'FL4-H' | 'Width' | |

### 2 Gradient Boosting Machine Learning (GBML) Suggested by Blasi et al.

To further investigate label-free quantification approaches using machine learning, we utilized the *gradient boosting machine learning* method, suggested by Blasi et al. [1]. The algorithm first partitions the data into training and testing sets, then conducts a cross-validation analysis on random partitions of the training data. Using the least squares boosting method in MATLAB's fitensemble routine, Blasi et al. have provided methods to conduct label-free quantification of the DNA content in fixed and live Jurkat cells. Predicting the DNA content based on darkfield and brightfield images from an imaging flow cytometer was deemed successful in their research. Nevertheless, their method failed to correctly identify the lipid content in our data (Fig. S4). The main reason for such a low prediction accuracy is that the best intrinsic cellular features are selected based solely on information collected from *fluorescently labeled* cells, which becomes problematic for the current study where non-label channels are affected by application of the lipid stain.

### 3 Multilayer Perceptron Neural Network (MLPNN)

In another approach, we utilized a multilayer perceptron neural network (MLPNN) which has multiple layers of logistic regression models with continuous nonlinearities. We chose an activation function $Tanh(x)$ in our MLPNN. The model we used is in the form

$$y_{\mathbf{k}}(\mathbf{x}, \mathbf{w}) = \sum_{j=0}^{H} w_{kj}^{(2)} Tanh(\sum_{i=0}^{N} w_{ji}^{(1)} x_i) \tag{1}$$

with $k$ outputs of $y_k$, where $H$ is the number of hidden units in a hidden layer, $N$ is number of input variables, $w_{kj}^{(2)}$'s are the weights in the second layer ($j = 1, ..., H$), $w_{ji}^{(1)}$'s are the weights in the first layer, and $x_i$ is the $i^{\text{th}}$ input variable ($i = 1, ..., N$). In the above equation, the bias parameters, $w_{j0}^{(1)}$ in the first layer, are included in the vector of weight parameters by adding an additional input $x_0 = 1$. A similar approach was taken for the second layer with respect to biases of the form $w_{k0}^{(2)}$. To optimize the weights and bias parameters, we used the scaled conjugate gradients back-propagation technique [2]. For the feature selection, the genetic algorithm was used to optimize the number of hidden units and feature combinations.
Figure S5 shows the results of this method on predicting the lipid content of the unlabeled cells.

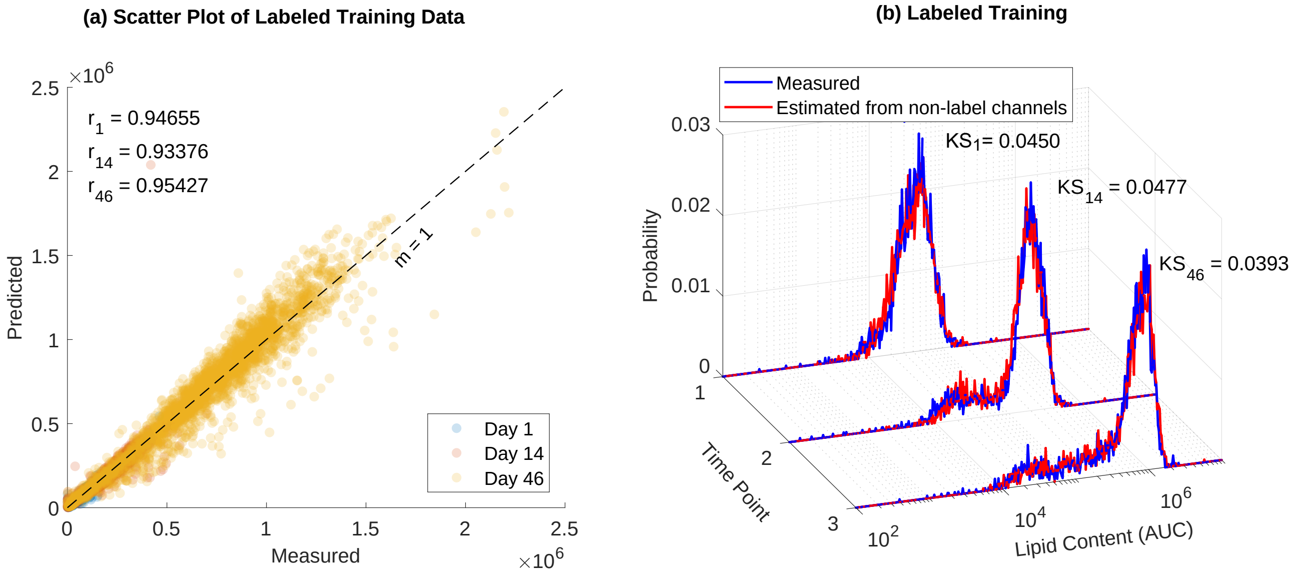

**Figure S. 1.** Preliminary regression analysis. (A) Correlations between measured and predicted values of lipid content for labeled training data. Pearsons correlation coefficients are shown for each time point. (B) Training the model to estimate the lipid content using non-label channels from labeled cell preparations. Measured in blue and predicted in red. Kolmogorov-Smirnov distances between the distributions are shown. Lipid content is shown in arbitrary units of concentration (AUC) in log scale.

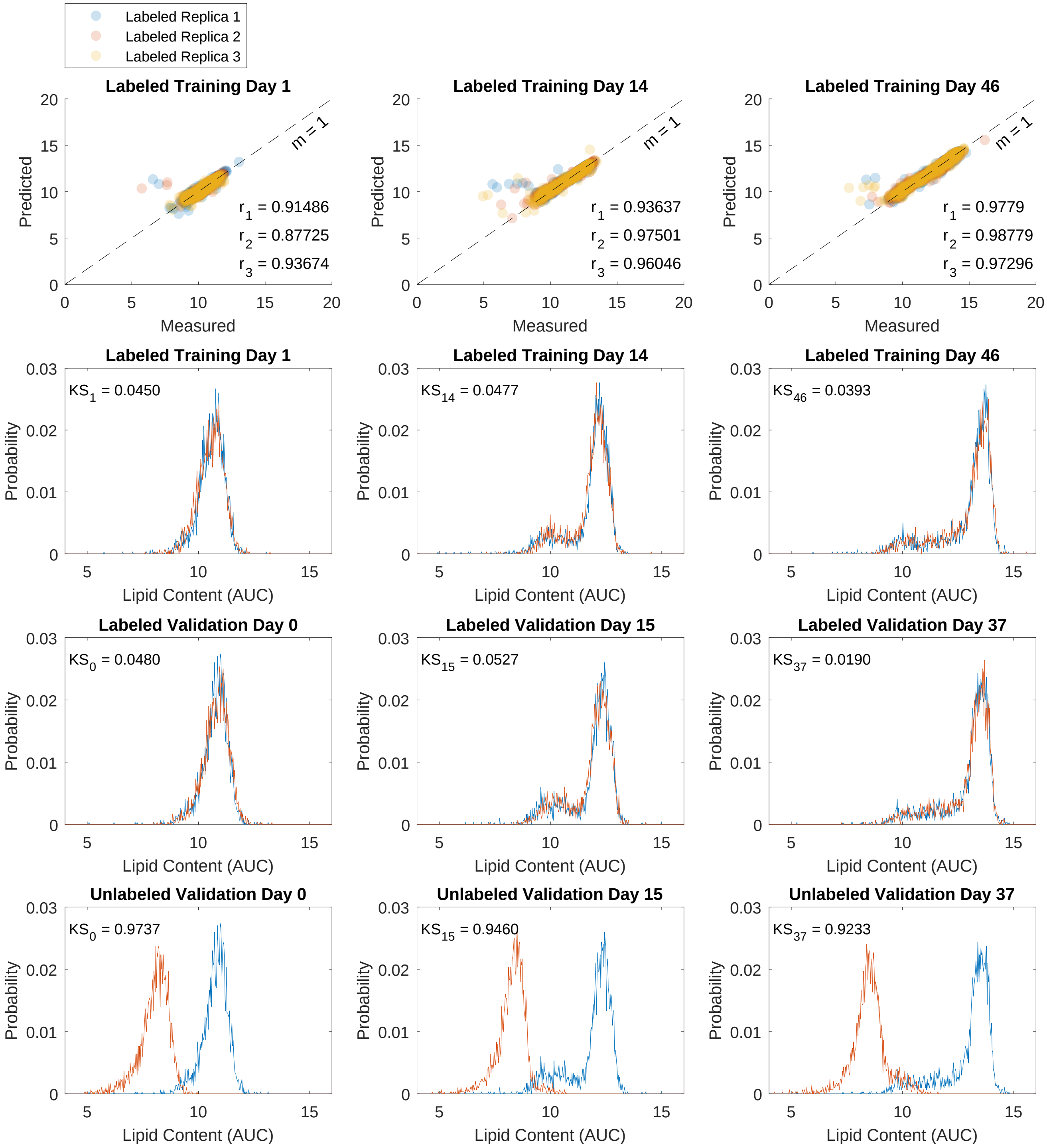

**Figure S. 2.** Preliminary regression analysis. The model works well for labeled data (rows 1, 2, and 3). However, it fails to predict the lipid content for unlabeled cells (row 4). The first row represents the correlation between measured and predicted values of the labeled training data. The 3 colors correspond to the 3 measurement replications at each day of FCM analysis (days 1, 14, and 46). Pearson's correlation coefficients are shown for each replication. For validation (rows 2, 3, and 4), we selected days 0, 15, and 37. The histograms show the results of prediction with this model for training (labeled cells) and validation (labeled and unlabeled cells) data. Measured histograms are in blue, predicted are in red. The KS distances between measured and predicted lipid content are shown on each plot. For all histograms, lipid content is shown in arbitrary units of concentration (AUC) in log scale.

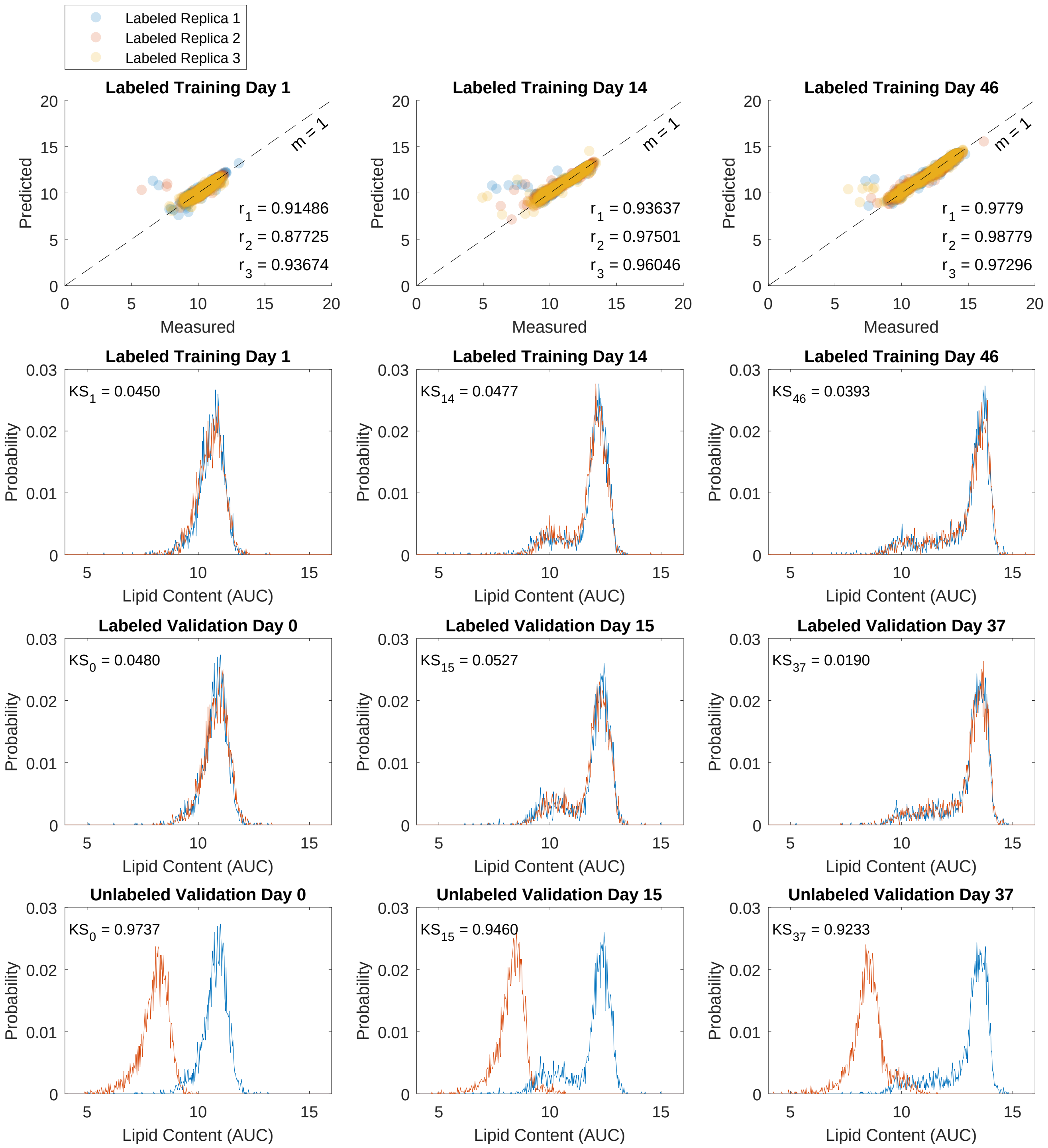

**Figure S. 3.** The quadratic regression model. The model works well for labeled data (rows 1, 2, and 3). However, it fails to predict the lipid content for unlabeled cells (row 4). The first row represents the correlation between measured and predicted values of the labeled training data. The 3 colors correspond to the 3 measurement replications at each day of FCM analysis (days 1, 14, and 46). Pearson's correlation coefficients are shown for each replication. For validation (rows 2, 3, and 4), we selected days 0, 15, and 37. The histograms show the results of prediction with this model for training (labeled cells) and validation (labeled and unlabeled cells) data. Measured histograms are in blue, predicted are in red. The KS distances between measured and predicted lipid content are shown on each plot. For all histograms, lipid content is shown in arbitrary units of concentration (AUC) in log scale.

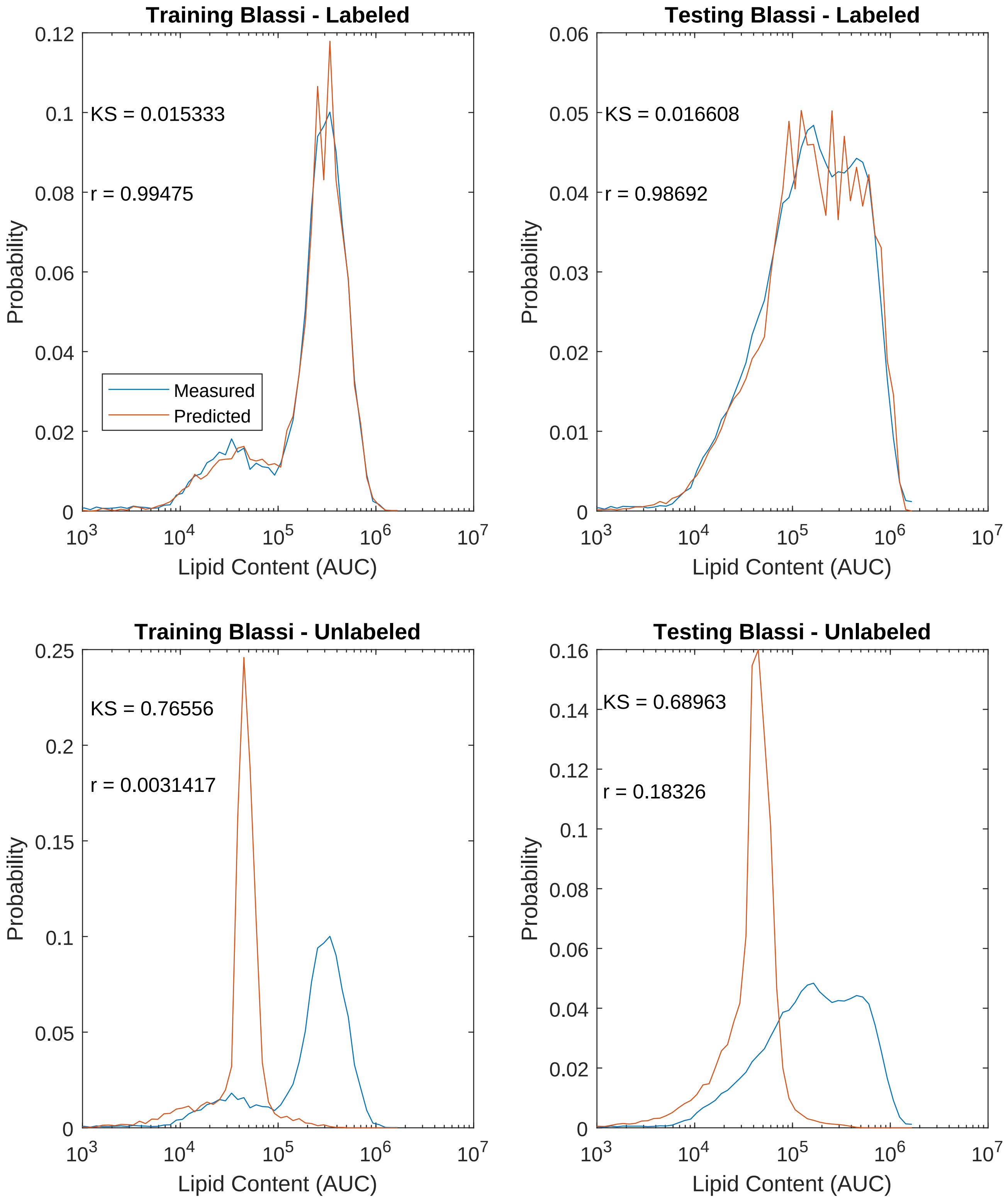

**Figure S. 4.** Testing the GBML on labeled and unlabeled data. The model works well for labeled cells, but fails drastically for unlabeled cells. Measured histograms are in blue, and predicted are in red. The KS distances between measured and predicted are shown. Pearson's correlation coefficients are shown. Lipid content is in arbitrary units of concentration (AUC).

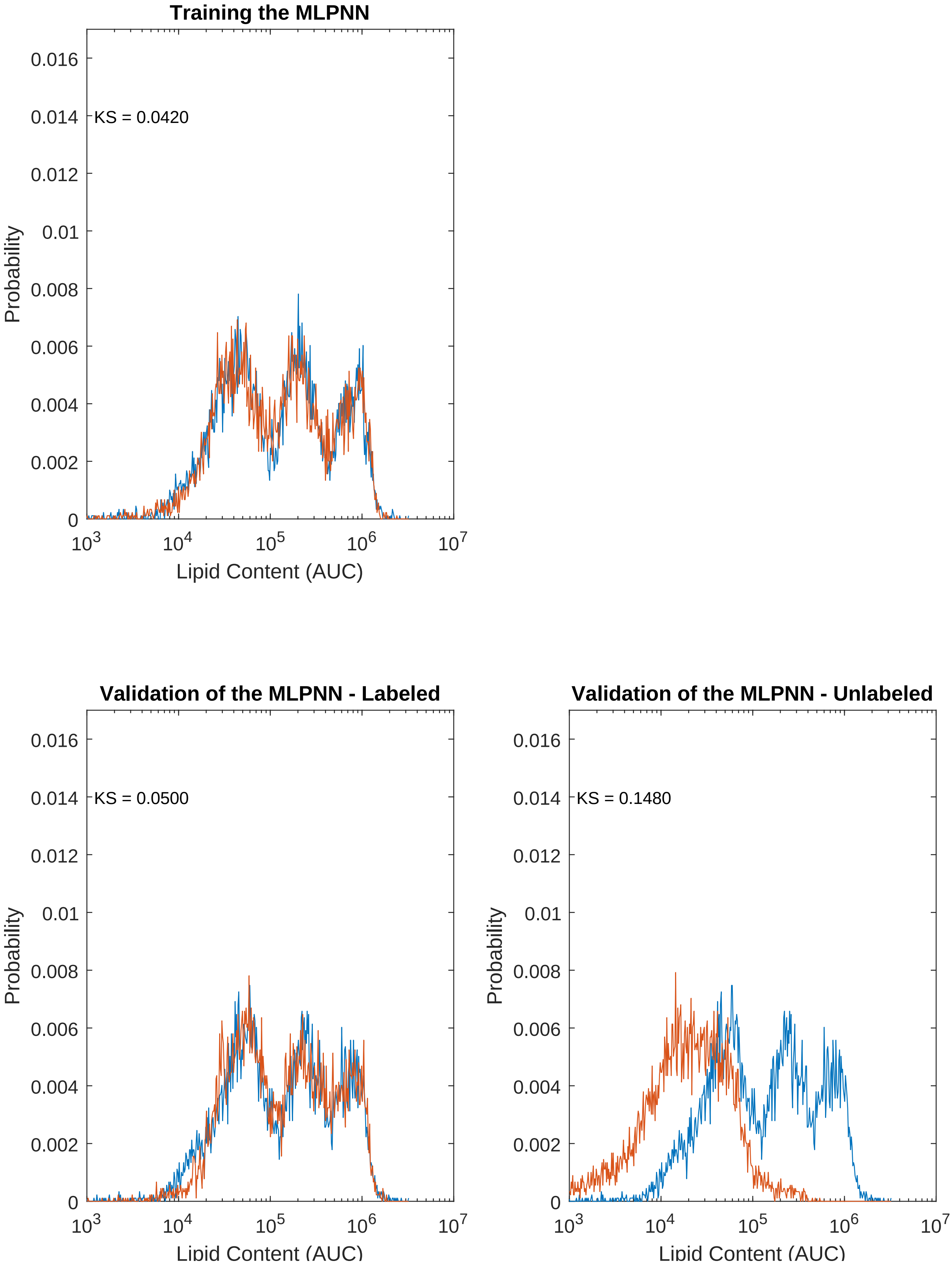

**Figure S. 5.** Results of using the MLPNN. Moderate prediction accuracy for both labeled and unlabeled cells. Measured histograms are in blue, predicted are in red. The KS distances between measured and predicted are shown. Lipid content is in arbitrary units of concentration (AUC).

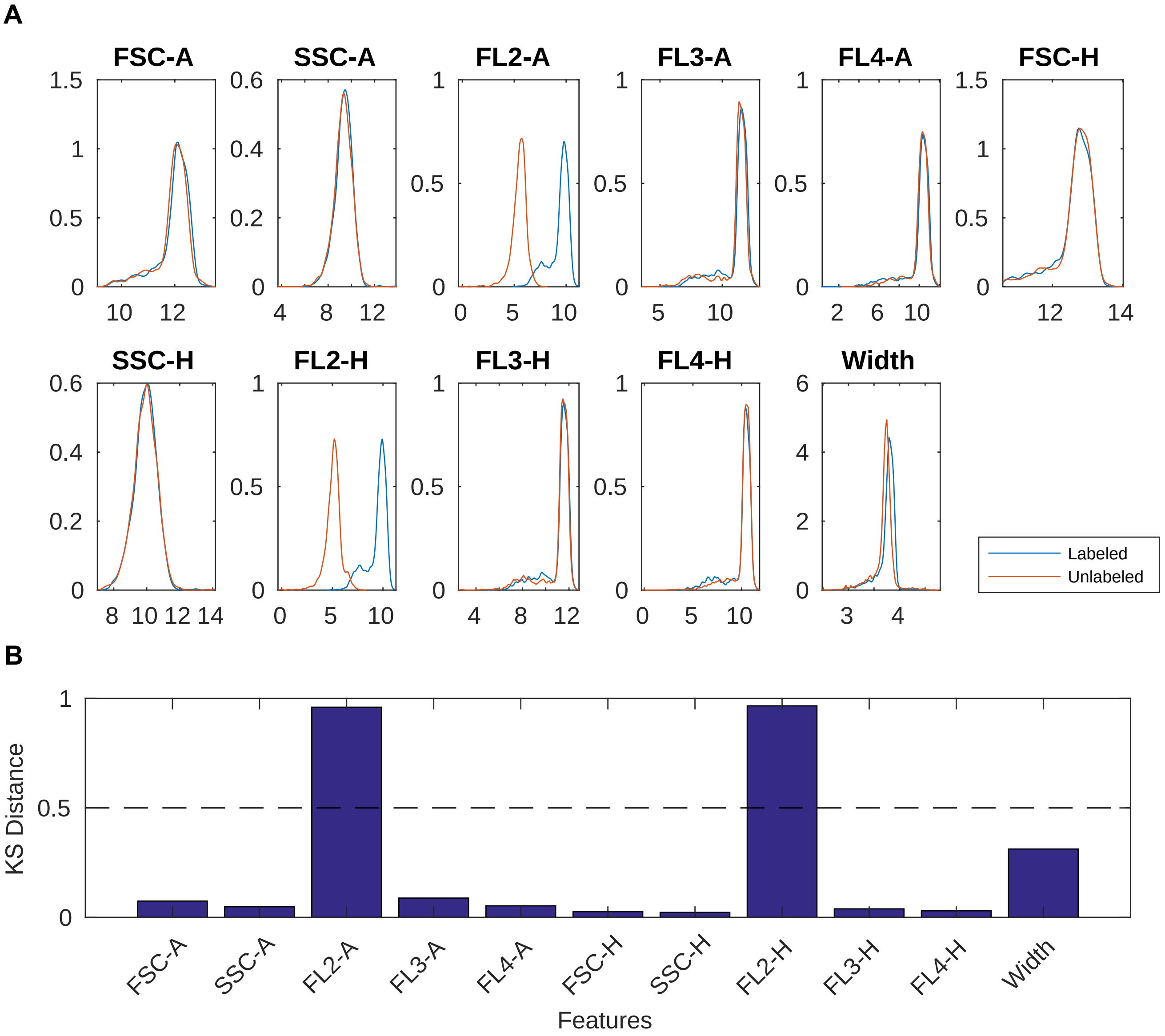

**Figure S. 6.** Comparison of the features with and without BODIPY stain and identification of the disrupted features. (A) Kernel densities of features for labeled and unlabeled cells, averaged over all times. Labeled cells are shown in blue, and unlabeled cells are in red. (B) KS distance between labeled and unlabeled features distributions. FL2-A and FL2-H features show clear dependence on the BODIPY stain. Horizontal line denotes threshold used to remove corrupted features.

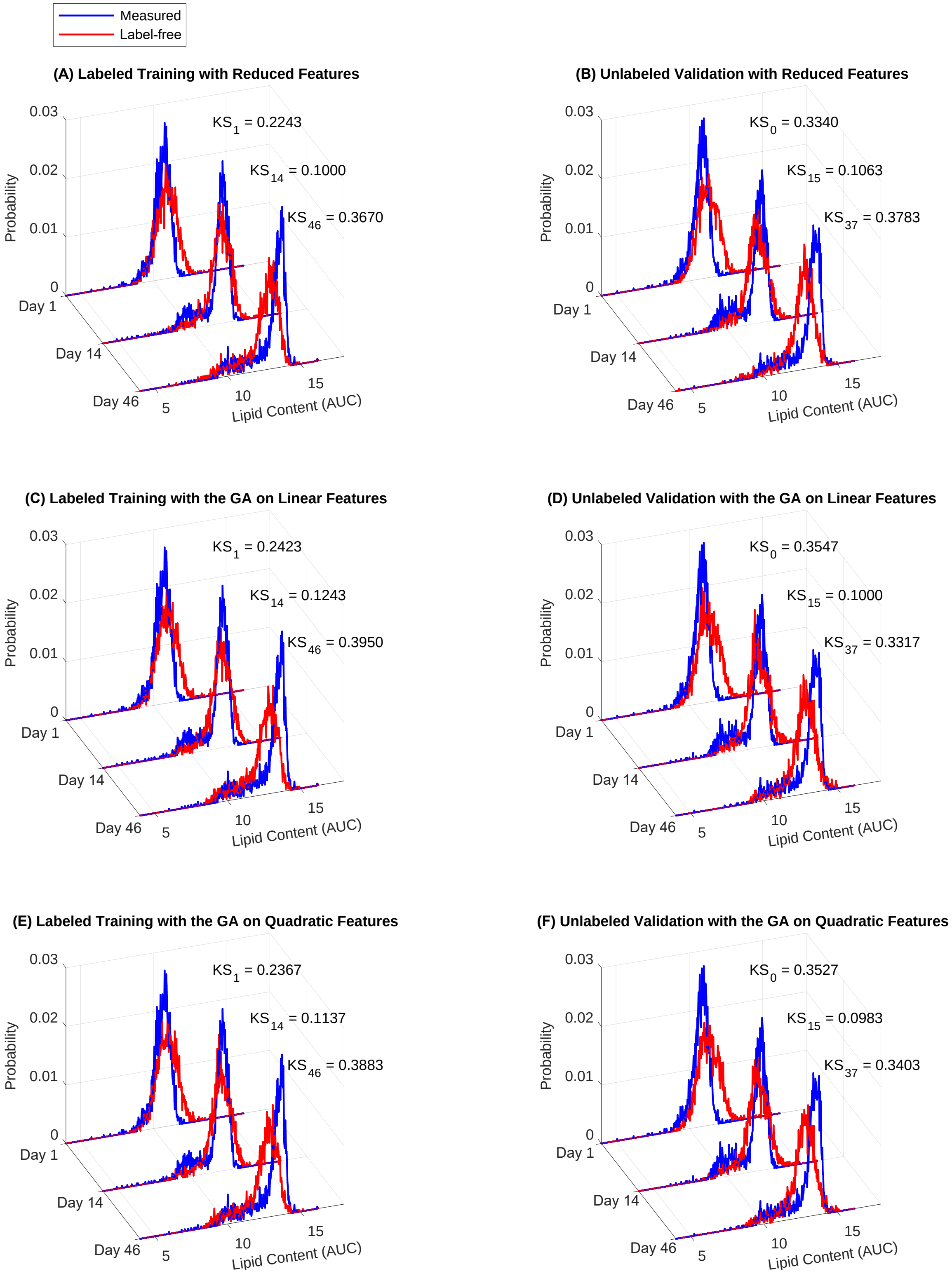

**Figure S. 7.** Regression results after various approaches to feature selection. (A) Training on reduced features. (B) Validation of the model in (A) on unlabeled cells. (C) Training based on the features selected by the genetic algorithm. (D) Validation of the model in (C) on unlabeled cells. (E) Training based on the features selected by the genetic algorithm on quadratic features and interactions. (F) Validation of the model in (E) on unlabeled cells. For all cases, measured values are shown in blue and predicted in red. Kolmogorov-Smirnov distances between distributions are shown. Training data correspond to days 1, 14, and 46; validation data correspond to days 0, 15, and 37. Lipid content is shown in arbitrary units of concentration (AUC) in log scale.

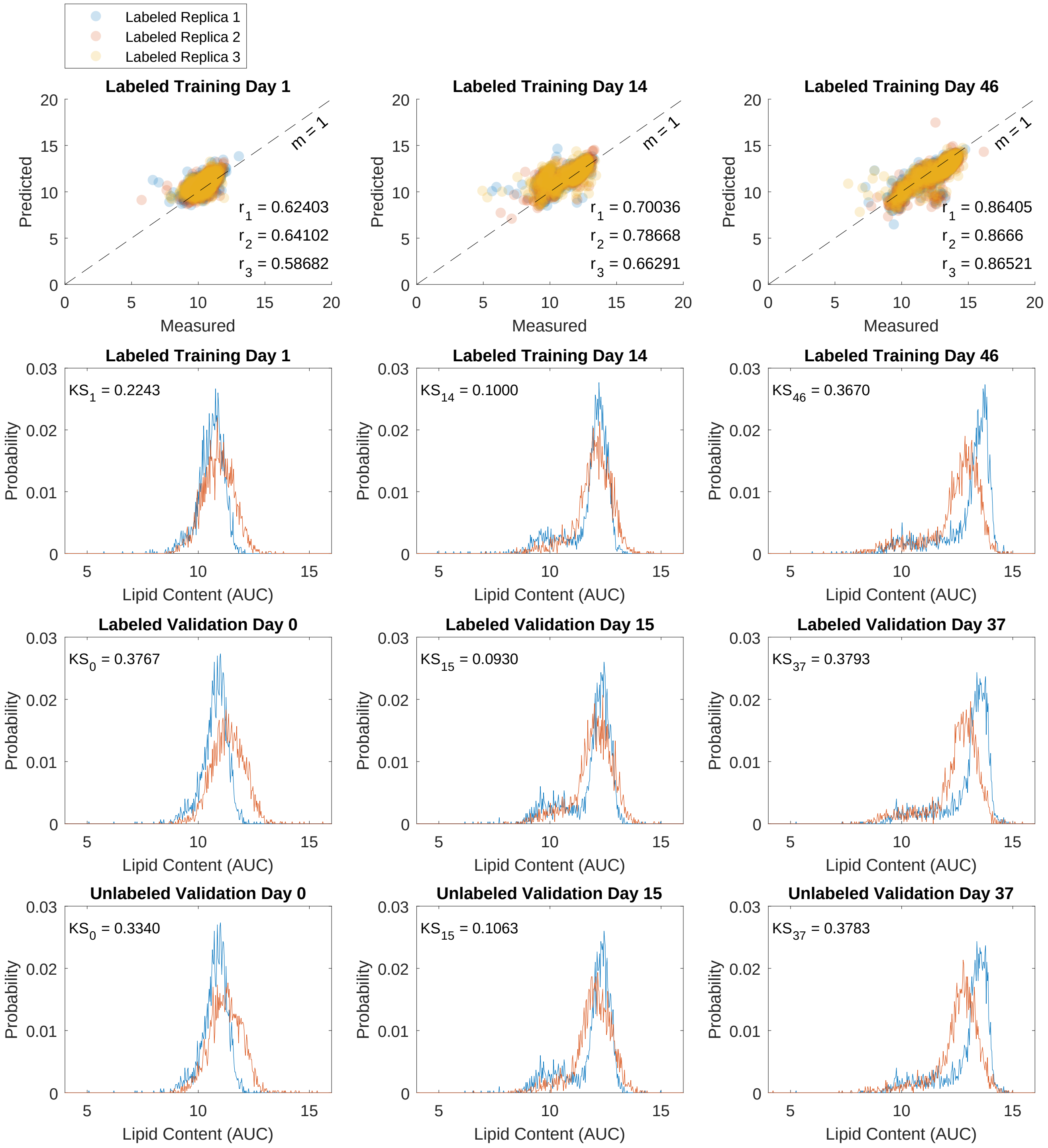

**Figure S. 8.** Linear regression on reduced features. Accuracy of the model is decreased for labeled data (rows 1, 2, and 3) due to removing the disrupted features. However, we see an important improvement in predicting the lipid content for the unlabeled cells (row 4). The first row represents the correlation between measured and predicted values of the labeled training data. The 3 colors correspond to the 3 measurement replications at each day of FCM analysis (days 1, 14, and 46). Pearson's correlation coefficients are shown for each replication. For validation (rows 2, 3, and 4), we selected days 0, 15, and 37. The histograms show the results of prediction with this model for training (labeled cells) and validation (labeled and unlabeled cells) data. Measured histograms are in blue, predicted are in red. The KS distances between measured and predicted lipid content are shown on each plot. For all histograms, lipid content is shown in arbitrary units of concentration (AUC) in log scale.

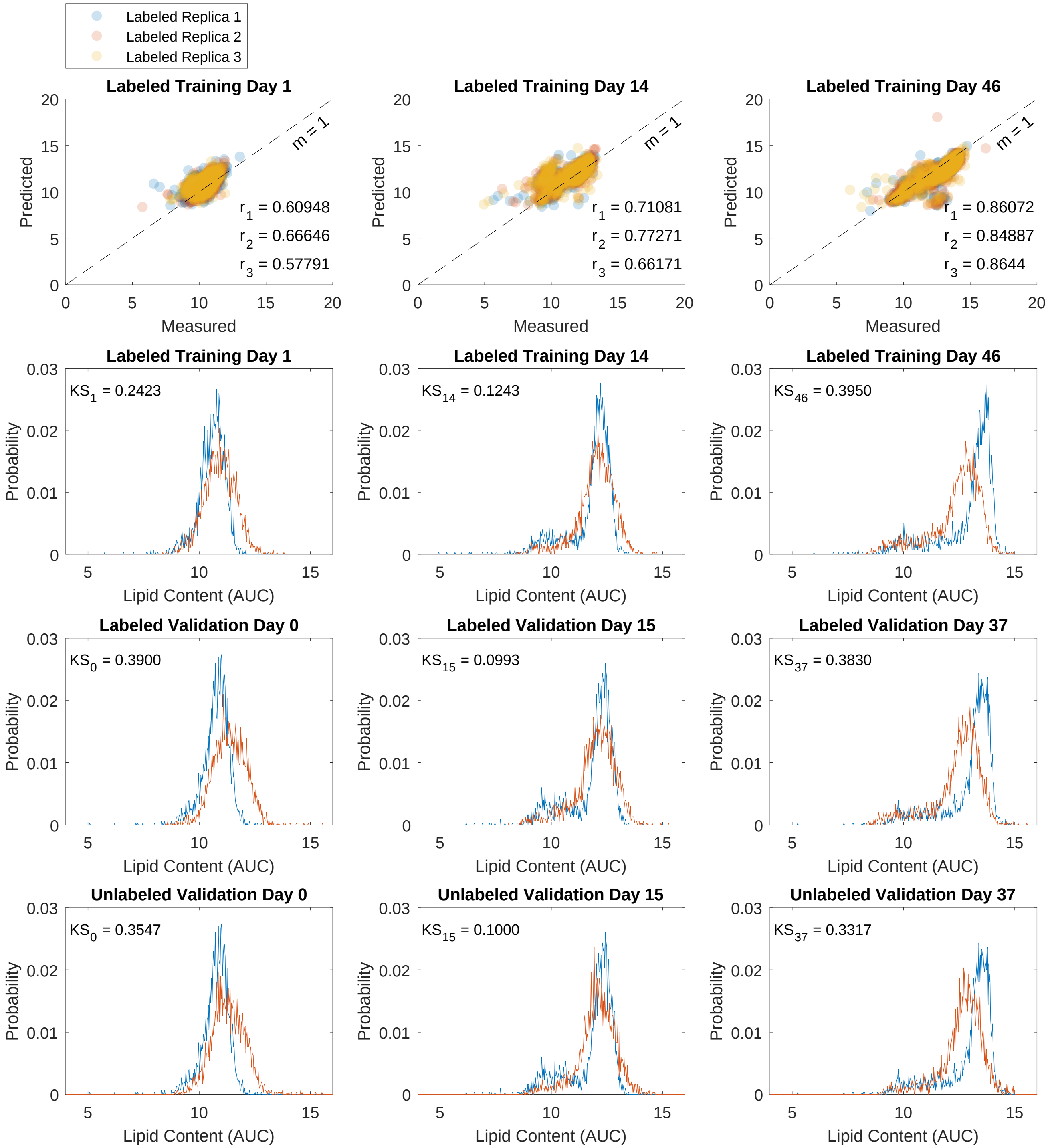

**Figure S. 9.** Regression analysis after performing automated feature selection by the genetic algorithm on linear features. Better prediction accuracy for the unlabeled cells. The first row represents the correlation between measured and predicted values of the labeled training data. The 3 colors correspond to the 3 measurement replications at each day of FCM analysis (days 1, 14, and 46). Pearson's correlation coefficients are shown for each replication. For validation (rows 2, 3, and 4), we selected days 0, 15, and 37. The histograms show the results of prediction with this model for training (labeled cells) and validation (labeled and unlabeled cells) data. Measured histograms are in blue, predicted are in red. The KS distances between measured and predicted lipid content are shown on each plot. For all histograms, lipid content is shown in arbitrary units of concentration (AUC) in log scale.

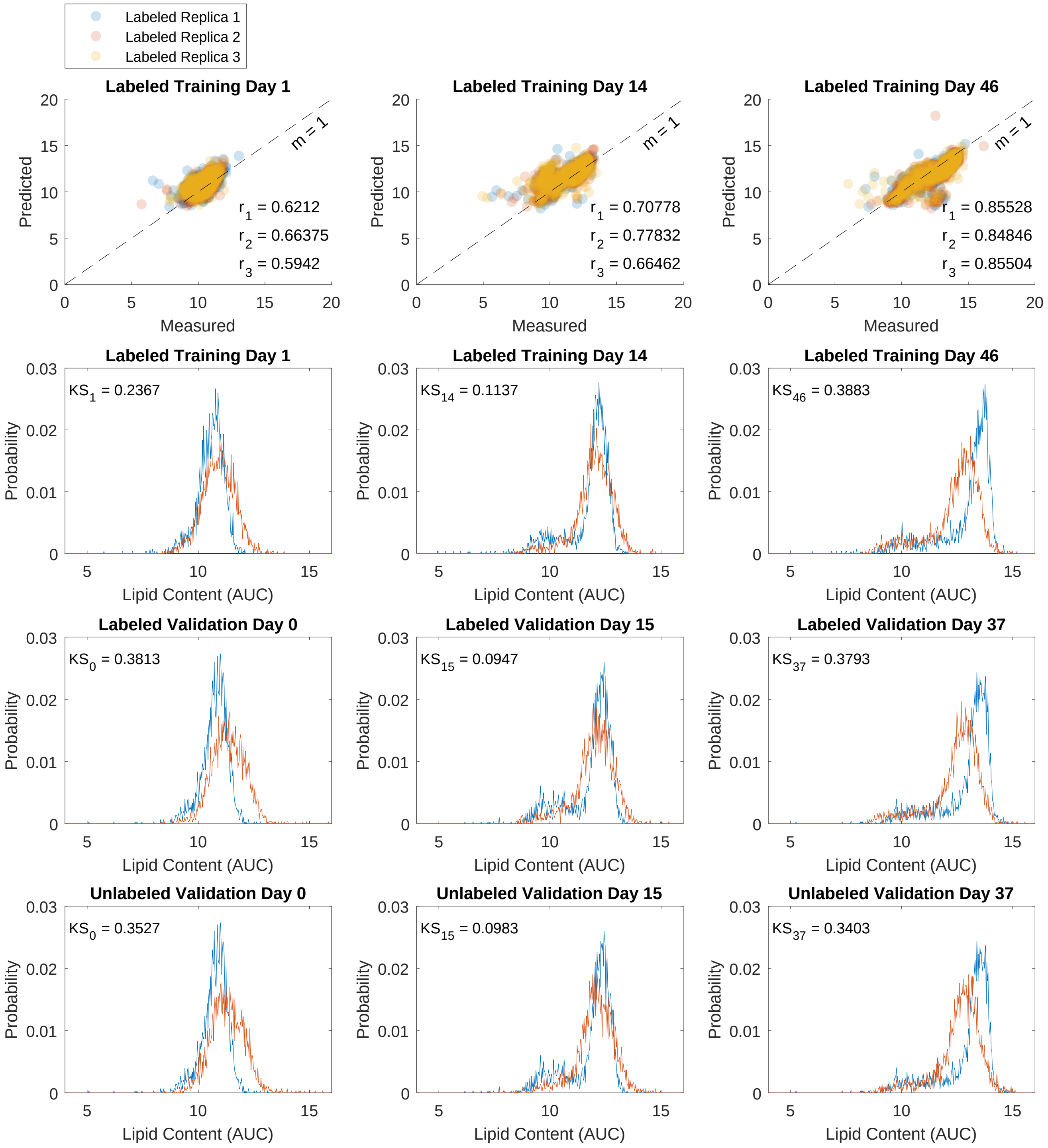

**Figure S. 10.** Regression analysis after performing automated feature selection by the genetic algorithm on quadratic features. Slight improvement is observed for prediction of the unlabeled cells' lipid content. The first row represents the correlation between measured and predicted values of the labeled training data. The 3 colors correspond to the 3 measurement replications at each day of FCM analysis (days 1, 14, and 46). Pearson's correlation coefficients are shown for each replication. For validation (rows 2, 3, and 4), we selected days 0, 15, and 37. The histograms show the results of prediction with this model for training (labeled cells) and validation (labeled and unlabeled cells) data. Measured histograms are in blue, predicted are in red. The KS distances between measured and predicted lipid content are shown on each plot. For all histograms, lipid content is shown in arbitrary units of concentration (AUC) in log scale.

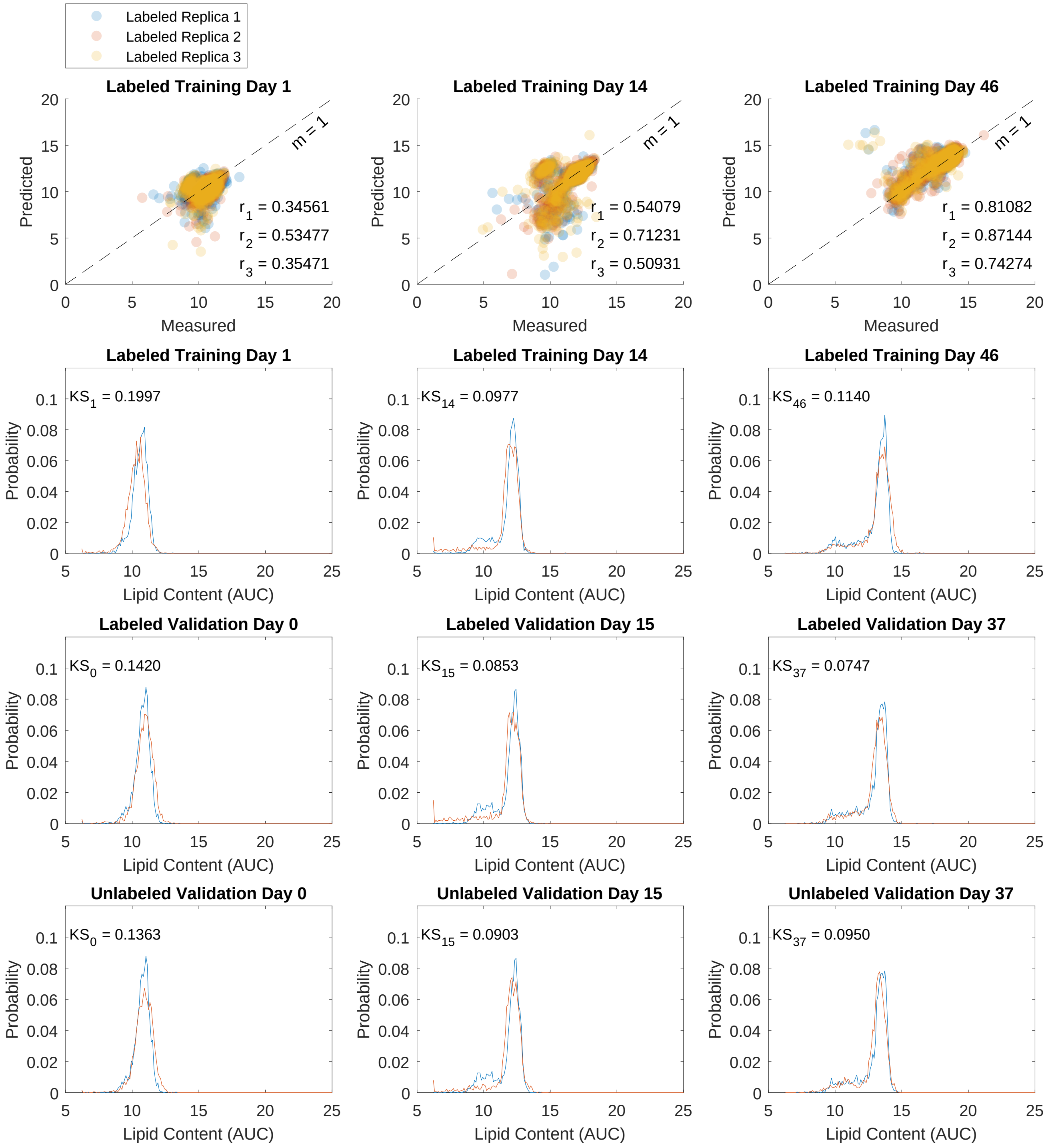

**Figure S. 11.** Our optimized label-free quantification strategy based on weighted models. The weights applied to the 3 trained models were estimated (using a secondary regression analysis) by measured tests statistics of the unlabeled features. The model was able to predict the lipid content of unlabeled cells with a remarkable high accuracy. The first row represents the correlation between measured and predicted values of the labeled training data. The 3 colors correspond to the 3 measurement replications at each day of FCM analysis (days 1, 14, and 46). Pearson's correlation coefficients are shown for each replication. For validation (rows 2, 3, and 4), we selected days 0, 15, and 37. The histograms show the results of prediction with this model for training (labeled cells) and validation (labeled and unlabeled cells) data. Measured histograms are in blue, predicted are in red. The KS distances between measured and predicted lipid content are shown on each plot. For all histograms, lipid content is shown in arbitrary units of concentration (AUC) in log scale.

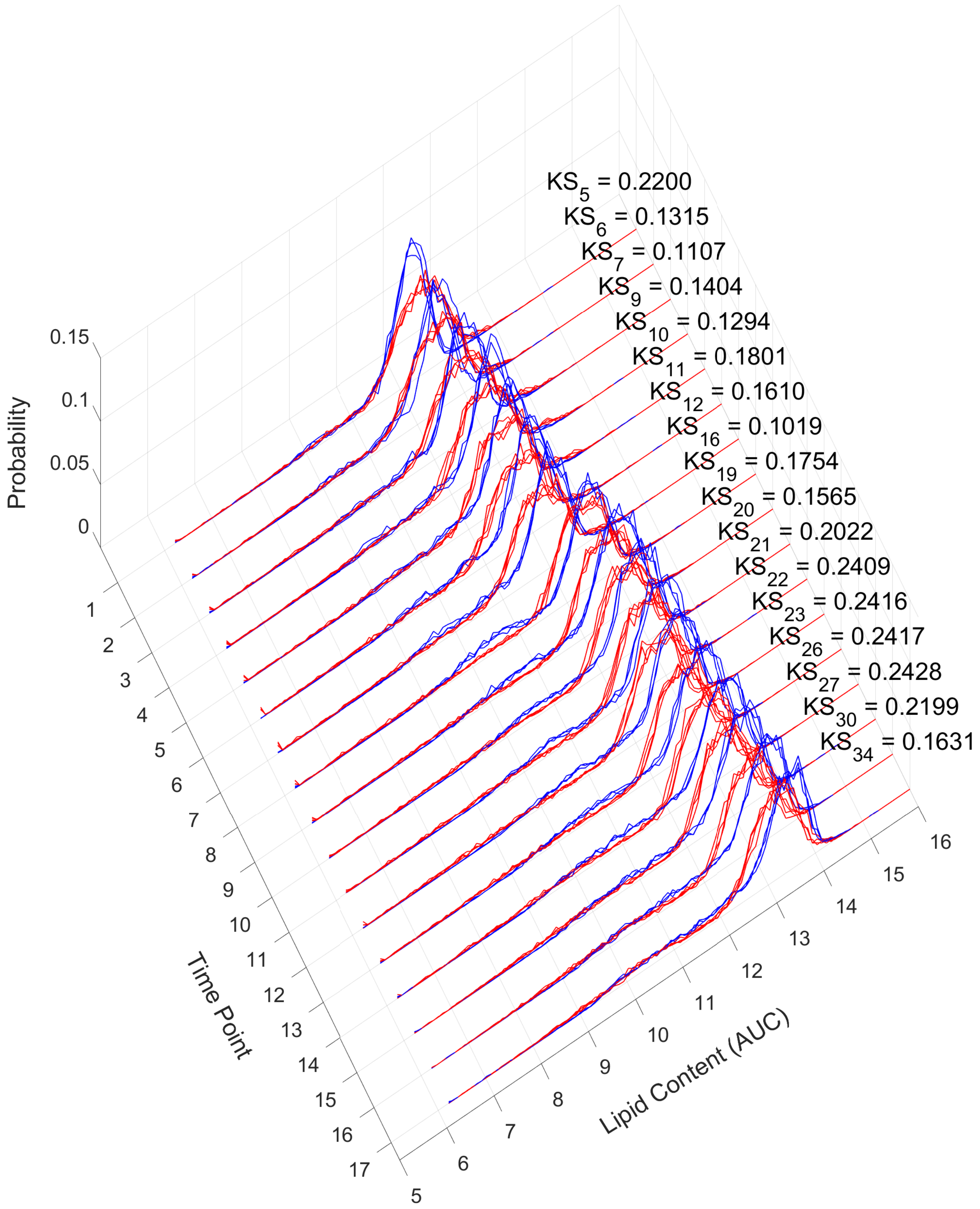

**Figure S. 12.** Testing the weighted model on all 17 testing time points. The data were not seen previously by the model. Measured histograms are in blue, predicted are in red. The average KS distances between measured and predicted are shown. Lipid content is shown in arbitrary units of concentration (AUC) in log scale.

**Table S. 1.** Feature selection by the genetic algorithm. We used 11 features in our analysis of the flow cytometric measurements. Features FL1-A and FL1-H were not included as they correspond to the main BODIPY fluorescence channel (targets to be predicted) and hence, they are not considered in the genetic algorithm.

**Table S. 1. 1.** The regular approach. Application of the genetic algorithm on linear features (selected features are shown in green). Regression coefficient values for each selected feature are shown.

| | |
|---|---|
| 'FSC-A' | 6.3395 |
| 'SSC-A' | 0.0548 |
| 'FL2-A' | 0 |
| 'FL3-A' | -0.1703 |
| 'FL4-A' | 0 |
| 'FSC-H' | -2.9008 |
| 'SSC-H' | 0 |
| 'FL2-H' | 0 |
| 'FL3-H' | 0 |
| 'FL4-H' | 0 |
| 'Width' | -6.8632 |

**Table S. 1. 2.** The regular approach. Application of the genetic algorithm on quadratic features (selected features are shown in green). Regression coefficient values for each selected feature are shown.

| | | | Feature | | | | | | | | | | |
|---|---|---|---|---|---|---|---|---|---|---|---|---|---|
| | | | 'FSC-A' | 'SSC-A' | 'FL2-A' | 'FL3-A' | 'FL4-A' | 'FSC-H' | 'SSC-H' | 'FL2-H' | 'FL3-H' | 'FL4-H' | 'Width' |
| Feature | 'FSC-A' | 0 | 0 | 0 | 0 | 0 | 0 | 0 | 0 | 0 | 0 | 6.7021 | 0 |
| | 'SSC-A' | 0 | | 0 | 0 | 0 | 0 | 0 | 0 | 0 | 0 | 0 | 0 |
| | 'FL2-A' | 0 | | | 0 | 0 | 0 | 0 | 0 | 0 | 0 | 0 | 0 |
| | 'FL3-A' | 0 | | | | 0 | 0 | 0 | 0 | 0 | 0 | 0 | 0 |
| | 'FL4-A' | 0 | | | | | 0 | 0 | 0 | 0 | 0 | 0 | 0 |
| | 'FSC-H' | 0 | | | | | | -1.5845 | 0 | 0 | 0 | 0 | 0 |
| | 'SSC-H' | 0 | | | | | | | 0 | 0 | 0 | 0 | 0 |
| | 'FL2-H' | 0 | | | | | | | | 0 | 0 | 0 | 0 |
| | 'FL3-H' | 0 | | | | | | | | | 0 | 0 | 0 |
| | 'FL4-H' | 0 | | | | | | | | | | 0 | -6.9141 |
| | 'Width' | 0 | | | | | | | | | | | 0 |

**Table S. 1. 3.** Our proposed strategy (weighted model). Application of the genetic algorithm on linear features (selected features are shown in green). The 3 trained models will have different weights. The genetic algorithm is performed to select the most informative features of each model separately. Regression coefficient values for each selected feature are shown.

| | | | |
|---|---|---|---|
| 'FSC-A' | 0 | 0 | 0 |
| 'SSC-A' | 0.3116 | 0 | 0.5483 |
| 'FL2-A' | 0 | 0 | 0 |
| 'FL3-A' | 0 | 0 | -0.0983 |
| 'FL4-A' | -0.0754 | 1.1708 | -0.1790 |
| 'FSC-H' | 0 | 0 | 0 |
| 'SSC-H' | 0 | 0 | 0 |
| 'FL2-H' | 0 | 0 | 0 |
| 'FL3-H' | 0 | 0 | 0.9724 |
| 'FL4-H' | 0.9466 | 0 | 0 |
| 'Width' | 0 | 0 | 0 |

**Table S. 1.** (continued)

**Table S. 1. 4.** Our proposed strategy (weighted model). Application of the genetic algorithm on quadratic features (selected features are shown in green). The genetic algorithm is applied to the 3 models M1, M2, and M3 separately.

Model 1

| | | | Feature (Model 1) | | | | | | | | | | |
|---|---|---|---|---|---|---|---|---|---|---|---|---|---|
| | | | 'FSC-A' | 'SSC-A' | 'FL2-A' | 'FL3-A' | 'FL4-A' | 'FSC-H' | 'SSC-H' | 'FL2-H' | 'FL3-H' | 'FL4-H' | 'Width' |
| Feature (Model 1) | 'FSC-A' | 0 | 0 | 0 | 0 | 0 | 0 | 0 | 0 | 0 | 0 | 0 | 0 |
| | 'SSC-A' | 0 | | 0 | 0 | -0.5060 | 0 | 0 | 0 | 0 | 0.9099 | 0 | 0 |
| | 'FL2-A' | 0 | | | 0 | 0 | 0 | 0 | 0 | 0 | 0 | 0 | 0 |
| | 'FL3-A' | 0 | | | | 0 | 0 | 0 | 0 | 0 | 0 | 0 | 0 |
| | 'FL4-A' | 0 | | | | | 0.1716 | 0 | 0 | 0 | 0 | 0 | 0 |
| | 'FSC-H' | 0 | | | | | | 0 | 0 | 0 | 0 | 0 | 0 |
| | 'SSC-H' | 0 | | | | | | | 0 | 0 | 0 | 0 | 0 |
| | 'FL2-H' | 0 | | | | | | | | 0 | 0 | 0 | 0 |
| | 'FL3-H' | 0 | | | | | | | | | 0 | 0 | 0 |
| | 'FL4-H' | 0 | | | | | | | | | | 0 | 0 |
| | 'Width' | 0 | | | | | | | | | | | 0 |

Model 2

| | | | Feature (Model 2) | | | | | | | | | | |
|---|---|---|---|---|---|---|---|---|---|---|---|---|---|
| | | | 'FSC-A' | 'SSC-A' | 'FL2-A' | 'FL3-A' | 'FL4-A' | 'FSC-H' | 'SSC-H' | 'FL2-H' | 'FL3-H' | 'FL4-H' | 'Width' |
| Feature (Model 2) | 'FSC-A' | 0 | 0 | 0 | 0 | 0 | 0 | 0 | 0 | 0 | 0 | 0 | 0 |
| | 'SSC-A' | 0 | | 0 | 0 | 0 | 0 | 0 | 0 | 0 | 0 | 0 | 0 |
| | 'FL2-A' | 0 | | | 0 | 0 | 0 | 0 | 0 | 0 | 0 | 0 | 0 |
| | 'FL3-A' | 0 | | | | 0 | 0 | 0 | 0 | 0 | 0 | 0 | 0 |
| | 'FL4-A' | 0 | | | | | 0.5854 | 0 | 0 | 0 | 0 | 0 | 0 |
| | 'FSC-H' | 0 | | | | | | 0 | 0 | 0 | 0 | 0 | 0 |
| | 'SSC-H' | 0 | | | | | | | 0 | 0 | 0 | 0 | 0 |
| | 'FL2-H' | 0 | | | | | | | | 0 | 0 | 0 | 0 |
| | 'FL3-H' | 0 | | | | | | | | | 0 | 0 | 0 |
| | 'FL4-H' | 0 | | | | | | | | | | 0 | 0 |
| | 'Width' | 0 | | | | | | | | | | | 0 |

Model 3

| | | | Feature (Model 3) | | | | | | | | | | |
|---|---|---|---|---|---|---|---|---|---|---|---|---|---|
| | | | 'FSC-A' | 'SSC-A' | 'FL2-A' | 'FL3-A' | 'FL4-A' | 'FSC-H' | 'SSC-H' | 'FL2-H' | 'FL3-H' | 'FL4-H' | 'Width' |
| Feature (Model 3) | 'FSC-A' | 0 | 0 | 0 | 0 | 0 | 0 | 0 | 0 | 0 | 0 | 0 | 0 |
| | 'SSC-A' | 0 | | 0 | 0 | 0 | 0 | 0 | 0.3494 | 0 | 0 | 0 | 0 |
| | 'FL2-A' | 0 | | | 0 | 0 | 0 | 0 | 0 | 0 | 0 | 0 | 0 |
| | 'FL3-A' | 0 | | | | -0.0629 | 0 | 0 | 0 | 0 | 0 | 0 | 0 |
| | 'FL4-A' | 0 | | | | | 0 | 0 | -0.1393 | 0 | 0 | 0 | 0 |
| | 'FSC-H' | 0 | | | | | | 0 | 0 | 0 | 0 | 0 | 0 |
| | 'SSC-H' | 0 | | | | | | | 0 | 0 | 0 | 0 | 0 |
| | 'FL2-H' | 0 | | | | | | | | 0 | 0 | 0 | 0 |
| | 'FL3-H' | 0 | | | | | | | | | 0.4719 | 0 | 0 |
| | 'FL4-H' | 0 | | | | | | | | | | 0 | 0 |
| | 'Width' | 0 | | | | | | | | | | | 0 |

**Table S. 2.** After performing a secondary regression analysis, our strategy yielded a weight quotient (containing information related to the means and standard deviations of the label-free measurements – see Methods section) used to calculate each corresponding weight ($\alpha_1$, $\alpha_2$, and $\alpha_3$). As a result of using the genetic algorithm, the most important columns of the test statistic (or rows of the weight quotient) are selected. Selected information by the genetic algorithm are indicated by green boxes.

**Table S. 2. 1.** The means of the features, their quadratic values, and their pairwise interactions.

| | | | Feature Means (Model 1) | | | | | | | | | | | | |
|---|---|---|---|---|---|---|---|---|---|---|---|---|---|---|---|
| | | | 'FSC-A' | 'SSC-A' | 'FL1-A' | 'FL2-A' | 'FL3-A' | 'FL4-A' | 'FSC-H' | 'SSC-H' | 'FL1-H' | 'FL2-H' | 'FL3-H' | 'FL4-H' | 'Width' |
| Feature Means for $\alpha_1$ | 'FSC-A' | 0 | 0 | 0 | 0 | 0 | 0 | 0 | 0 | 0 | 0 | 0 | 0 | 0 | 0 |
| | 'SSC-A' | 0 | | -0.0002 | 0 | 0 | 0 | 0 | 0 | 0 | 0 | 0 | 0 | 0 | 0 |
| | 'FL1-A' | 0 | | | 0 | 0 | 0 | 0 | 0 | 0 | 0 | 0 | 0 | 0 | 0 |
| | 'FL2-A' | 0 | | | | 0 | 0 | 0 | 0 | 0 | 0 | 0 | 0 | 0 | 0 |
| | 'FL3-A' | 0 | | | | | 0 | 0 | 0 | 0 | 0 | 0 | 0 | 0 | 0 |
| | 'FL4-A' | 0 | | | | | | 0.7151 | 0 | 0 | 0 | 0 | 0 | 0 | 0 |
| | 'FSC-H' | 0 | | | | | | | 0.1017 | 0 | 0 | 0 | 0 | 0 | 0 |
| | 'SSC-H' | 0 | | | | | | | | 0 | 0 | 0 | 0 | 0 | 0 |
| | 'FL1-H' | 0 | | | | | | | | | -1.6985 | 0 | 0 | 0 | 0 |
| | 'FL2-H' | 0 | | | | | | | | | | 0 | 0 | 0 | 0 |
| | 'FL3-H' | 0 | | | | | | | | | | | 0 | 0 | 0 |
| | 'FL4-H' | 0 | | | | | | | | | | | | 0 | 0 |
| | 'Width' | 0 | | | | | | | | | | | | | 0 |

| | | | Feature Means (Model 2) | | | | | | | | | | | | |
|---|---|---|---|---|---|---|---|---|---|---|---|---|---|---|---|
| | | | 'FSC-A' | 'SSC-A' | 'FL1-A' | 'FL2-A' | 'FL3-A' | 'FL4-A' | 'FSC-H' | 'SSC-H' | 'FL1-H' | 'FL2-H' | 'FL3-H' | 'FL4-H' | 'Width' |
| Feature Means for $\alpha_2$ | 'FSC-A' | 0 | 0 | 0 | 0 | 0 | 0 | 0 | 0 | 0 | 0 | 0 | 0 | 0 | 0 |
| | 'SSC-A' | 0 | | -0.5351 | 0 | 0 | 0 | 0 | 0 | 0 | 0 | 0 | 0 | 0 | 0 |
| | 'FL1-A' | 0 | | | 0 | 0 | 0 | 0 | 0 | 0 | 0 | 0 | 0 | 0 | 0 |
| | 'FL2-A' | 0 | | | | 0 | 0 | 0 | 0 | 0 | 0 | 0 | 0 | 0 | 0 |
| | 'FL3-A' | 0 | | | | | 0 | 0 | 0 | 0 | 0 | 0 | 0 | 0 | 0 |
| | 'FL4-A' | 0 | | | | | | -0.2651 | 0 | 0 | 0 | 0 | 0 | 0 | 0 |
| | 'FSC-H' | 0 | | | | | | | 0.2427 | 0 | 0 | 0 | 0 | 0 | 0 |
| | 'SSC-H' | 0 | | | | | | | | 0 | 0 | 0 | 0 | 0 | 0 |
| | 'FL1-H' | 0 | | | | | | | | | 1.2247 | 0 | 0 | 0 | 0 |
| | 'FL2-H' | 0 | | | | | | | | | | 0 | 0 | 0 | 0 |
| | 'FL3-H' | 0 | | | | | | | | | | | 0 | 0 | 0 |
| | 'FL4-H' | 0 | | | | | | | | | | | | 0 | 0 |
| | 'Width' | 0 | | | | | | | | | | | | | 0 |

| | | | Feature Means (Model 3) | | | | | | | | | | | | |
|---|---|---|---|---|---|---|---|---|---|---|---|---|---|---|---|
| | | | 'FSC-A' | 'SSC-A' | 'FL1-A' | 'FL2-A' | 'FL3-A' | 'FL4-A' | 'FSC-H' | 'SSC-H' | 'FL1-H' | 'FL2-H' | 'FL3-H' | 'FL4-H' | 'Width' |
| Feature Means for $\alpha_3$ | 'FSC-A' | 0 | 0 | 0 | 0 | 0 | 0 | 0 | 0 | 0 | 0 | 0 | 0 | 0 | 0 |
| | 'SSC-A' | 0 | | 0.5597 | 0 | 0 | 0 | 0 | 0 | 0 | 0 | 0 | 0 | 0 | 0 |
| | 'FL1-A' | 0 | | | 0 | 0 | 0 | 0 | 0 | 0 | 0 | 0 | 0 | 0 | 0 |
| | 'FL2-A' | 0 | | | | 0 | 0 | 0 | 0 | 0 | 0 | 0 | 0 | 0 | 0 |
| | 'FL3-A' | 0 | | | | | 0 | 0 | 0 | 0 | 0 | 0 | 0 | 0 | 0 |
| | 'FL4-A' | 0 | | | | | | -0.4344 | 0 | 0 | 0 | 0 | 0 | 0 | 0 |
| | 'FSC-H' | 0 | | | | | | | -0.2852 | 0 | 0 | 0 | 0 | 0 | 0 |
| | 'SSC-H' | 0 | | | | | | | | 0 | 0 | 0 | 0 | 0 | 0 |
| | 'FL1-H' | 0 | | | | | | | | | 0.3706 | 0 | 0 | 0 | 0 |
| | 'FL2-H' | 0 | | | | | | | | | | 0 | 0 | 0 | 0 |
| | 'FL3-H' | 0 | | | | | | | | | | | 0 | 0 | 0 |
| | 'FL4-H' | 0 | | | | | | | | | | | | 0 | 0 |
| | 'Width' | 0 | | | | | | | | | | | | | 0 |

**Table S. 2.** (continued)

**Table S. 2. 2.** The standard deviations of the features, their quadratic values, and their pairwise interactions (model 1).

| | | | Feature Standard Deviations for $\alpha_1$ | | | | | | | | | | | | |
|---|---|---|---|---|---|---|---|---|---|---|---|---|---|---|---|
| | | | 'FSC-A' | 'SSC-A' | 'FL1-A' | 'FL2-A' | 'FL3-A' | 'FL4-A' | 'FSC-H' | 'SSC-H' | 'FL1-H' | 'FL2-H' | 'FL3-H' | 'FL4-H' | 'Width' |
| Feature Standard Deviations for $\alpha_1$ | 'FSC-A' | 0 | 0 | 0 | 0 | 0 | 0 | 0 | 0 | 0 | 0 | 0 | 0 | 0 | 0 |
| | 'SSC-A' | 0 | | 0 | 0 | 0 | 0 | 0 | 0 | 0 | 0 | 0 | 0 | 0 | 0 |
| | 'FL1-A' | 0 | | | 0 | 0 | 0 | 0 | 0 | 0 | 0 | 0 | 0 | 0 | 0 |
| | 'FL2-A' | 0 | | | | 0 | 0 | 0 | 0 | 0 | 0 | 0 | 0 | 0 | 0 |
| | 'FL3-A' | 0 | | | | | 0 | 0 | 0 | 0 | 0 | 0 | 0 | 0 | 0 |
| | 'FL4-A' | 0 | | | | | | 0 | 0 | 0 | 0 | 0 | 0 | 0 | 0 |
| | 'FSC-H' | 0 | | | | | | | 0 | 0 | 0 | 0 | 0 | 0 | 0 |
| | 'SSC-H' | 0 | | | | | | | | 0 | 0 | 0 | 0 | 0 | 0 |
| | 'FL1-H' | 0 | | | | | | | | | 0 | 0 | 0 | 0 | 0 |
| | 'FL2-H' | 0 | | | | | | | | | | 2.1244 | 0 | 0 | 0 |
| | 'FL3-H' | 0 | | | | | | | | | | | 0.4623 | 0 | 0 |
| | 'FL4-H' | 0 | | | | | | | | | | | | 0 | 0 |
| | 'Width' | 0 | | | | | | | | | | | | | 0 |

| | | | Feature Standard Deviations for $\alpha_2$ | | | | | | | | | | | | |
|---|---|---|---|---|---|---|---|---|---|---|---|---|---|---|---|
| | | | 'FSC-A' | 'SSC-A' | 'FL1-A' | 'FL2-A' | 'FL3-A' | 'FL4-A' | 'FSC-H' | 'SSC-H' | 'FL1-H' | 'FL2-H' | 'FL3-H' | 'FL4-H' | 'Width' |
| Feature Standard Deviations for $\alpha_2$ | 'FSC-A' | 0 | 0 | 0 | 0 | 0 | 0 | 0 | 0 | 0 | 0 | 0 | 0 | 0 | 0 |
| | 'SSC-A' | 0 | | 0 | 0 | 0 | 0 | 0 | 0 | 0 | 0 | 0 | 0 | 0 | 0 |
| | 'FL1-A' | 0 | | | 0 | 0 | 0 | 0 | 0 | 0 | 0 | 0 | 0 | 0 | 0 |
| | 'FL2-A' | 0 | | | | 0 | 0 | 0 | 0 | 0 | 0 | 0 | 0 | 0 | 0 |
| | 'FL3-A' | 0 | | | | | 0 | 0 | 0 | 0 | 0 | 0 | 0 | 0 | 0 |
| | 'FL4-A' | 0 | | | | | | 0 | 0 | 0 | 0 | 0 | 0 | 0 | 0 |
| | 'FSC-H' | 0 | | | | | | | 0 | 0 | 0 | 0 | 0 | 0 | 0 |
| | 'SSC-H' | 0 | | | | | | | | 0 | 0 | 0 | 0 | 0 | 0 |
| | 'FL1-H' | 0 | | | | | | | | | 0 | 0 | 0 | 0 | 0 |
| | 'FL2-H' | 0 | | | | | | | | | | -2.7642 | 0 | 0 | 0 |
| | 'FL3-H' | 0 | | | | | | | | | | | -0.6221 | 0 | 0 |
| | 'FL4-H' | 0 | | | | | | | | | | | | 0 | 0 |
| | 'Width' | 0 | | | | | | | | | | | | | 0 |

| | | | Feature Standard Deviations for $\alpha_3$ | | | | | | | | | | | | |
|---|---|---|---|---|---|---|---|---|---|---|---|---|---|---|---|
| | | | 'FSC-A' | 'SSC-A' | 'FL1-A' | 'FL2-A' | 'FL3-A' | 'FL4-A' | 'FSC-H' | 'SSC-H' | 'FL1-H' | 'FL2-H' | 'FL3-H' | 'FL4-H' | 'Width' |
| Feature Standard Deviations for $\alpha_3$ | 'FSC-A' | 0 | 0 | 0 | 0 | 0 | 0 | 0 | 0 | 0 | 0 | 0 | 0 | 0 | 0 |
| | 'SSC-A' | 0 | | 0 | 0 | 0 | 0 | 0 | 0 | 0 | 0 | 0 | 0 | 0 | 0 |
| | 'FL1-A' | 0 | | | 0 | 0 | 0 | 0 | 0 | 0 | 0 | 0 | 0 | 0 | 0 |
| | 'FL2-A' | 0 | | | | 0 | 0 | 0 | 0 | 0 | 0 | 0 | 0 | 0 | 0 |
| | 'FL3-A' | 0 | | | | | 0 | 0 | 0 | 0 | 0 | 0 | 0 | 0 | 0 |
| | 'FL4-A' | 0 | | | | | | 0 | 0 | 0 | 0 | 0 | 0 | 0 | 0 |
| | 'FSC-H' | 0 | | | | | | | 0 | 0 | 0 | 0 | 0 | 0 | 0 |
| | 'SSC-H' | 0 | | | | | | | | 0 | 0 | 0 | 0 | 0 | 0 |
| | 'FL1-H' | 0 | | | | | | | | | 0 | 0 | 0 | 0 | 0 |
| | 'FL2-H' | 0 | | | | | | | | | | 0.6488 | 0 | 0 | 0 |
| | 'FL3-H' | 0 | | | | | | | | | | | 0.1523 | 0 | 0 |
| | 'FL4-H' | 0 | | | | | | | | | | | | 0 | 0 |
| | 'Width' | 0 | | | | | | | | | | | | | 0 |
